# Long- and short-range bimodal signals mediate mate location and recognition in yellow fever mosquitoes

**DOI:** 10.1101/2021.04.27.441577

**Authors:** Elton Ko, Adam J. Blake, Chiara Lier, Stephen Takács, Gerhard Gries

## Abstract

Mate location and mate selection behavior in diurnally-active yellow fever mosquitoes, *Aedes aegypti*, take place in mating swarms but the mechanisms underlying swarm formation and long-range detection of females by males remain largely unexplored. In high-speed video recordings we show that incident light reflects off the wings of swarming males, and in behavioral experiments we demonstrate that swarm formation and mate recognition are mediated, in part, by these wingbeat light flashes and by wingbeat sound signals that operate at long and short range, respectively. To test for range-dependent effects of these signals, we presented ‘mating swarms’ in form of two paired 8-LED assemblies that were fitted with micro-speakers and placed either well separated in a large space or side-by-side in a small space. In the large but not the small space, the LED assembly flashing light at the wingbeat frequency of females (665 Hz), and emitting their wingbeat sound (665 Hz), attracted and prompted 5.8-times more alightings by males than the LED assembly emitting constant light and wingbeat sound. In the small space, the LED assembly flashing light and emitting wingbeat sound induced 5.0-times more alightings by males than the LED assembly flashing light without wingbeat sound. The attractiveness of light flash signals to males increased with increasing numbers of signals but did not vary according to their wavelengths (UV or blue). Females responded to light flash signals of males. As predicted by the sensory drive theory, light flashes had no signal function for crepuscular house mosquitoes, *Culex pipiens*.

## INTRODUCTION

Searching for a blood meal, female mosquitoes exploit multiple vertebrate host cues including CO_2_, body odor, moisture, as well as visual and heat contrast.^1^ To locate a host, female mosquitoes are guided by these chemical and physical cues in sequential and interactive processes.^2,3,4^ Exhaled in the breath of a potential host, CO_2_ context-dependently^5^ promotes host-seeking,^6,7^ elicits upwind flight toward the CO_2_ source,^8,9^ and enhances mosquito attraction to warmth.^10,11^ In addition to exhaled CO_2_, breath volatiles and numerous odorants emanating from bacteria on vertebrate skin^12—14^ guide host-foraging mosquitoes. The relative importance of host cues depends on the spatial scale, with some cues (thermal, skin odours, visual, moisture) being most important at close range.^4,15,16,17^

Nectar-foraging mosquitoes also exploit multimodal cues to locate floral resources.^18^ Females of the yellow fever mosquito, *Aedes aegypti*, and the Northern house mosquito, *Culex pipiens*, respond more strongly to a cue complex of tansy, *Tanacetum vulgare*, inflorescences, consisting of CO_2_, olfactory and visual cues, than to inflorescence odor alone.^18^ During floral foraging, floral odor likely acts as a long-range attractant, whereas visual cues are utilized at closer ranges.

While the multimodal sensory cues that guide foraging mosquitoes to host and nectar resources have been intensely studied, the mechanisms underlying mate location and recognition in mosquitoes are not fully understood. Many mosquito species form ‘mating swarms’ dominated by males^19^ that independently respond to ‘swarm marker’ objects in the environment^19^, such as trees, corn stalks, telephone poles^20^ or just black cards.^21^ In *Ae. aegypti*, the vertebrate host itself serves as the swarm marker.^22^ Swarming behavior exposes mosquitoes to predation^23^ and is energetically costly,^24^ but it expedites mate location which is challenging for species with widespread larval habitats.^19^

In the context of swarming, mosquitoes respond to acoustic, visual and pheromonal signals or cues from conspecifics.^25—27^ As shown for several species, swarming males recognize the wingbeat frequency of conspecific females that enter a swarm.^25,28,29^ The sound-receiving Johnston’s organ in the males’ antennae is attuned to the females’ wingbeat frequencies^25,29^ which are attractive to males.^30^ After successful coupling with the female, the male-female pair leaves the swarm to mate.^31^ In *Ae. aegypti*, courtship precedes coupling and entails harmonic convergence of both male and female wingbeat frequencies.^32^ Analogous behaviour has been reported in the elephant mosquito, *Toxorhynchites brevipalpis*,^33^ the southern house mosquito, *Culex quinquefasciatus*,^34^ and the African malaria mosquito, *Anopheles gambiae*.^35^

Males typically detect the wingbeat sound of females only at close range,^36^ indicating that physical mate location cues other than sound function at a longer range, as recently shown for several dipterans, including mosquitoes.^27,37^ Males of the common green bottle fly, *Lucilia sericata*, distinguish between the rates of light flashes reflected off the wings of in-flight female and male flies, and are most strongly attracted to flash frequencies (178 Hz) characteristic of young females.^37^ Similarly, 8-LED ‘mating swarm’ mimics of *Ae. aegypti* flashing white or blue light at the wing beat frequency of females (665 Hz) attract conspecific males.^27^ As thin-film reflectors,^37,38^ sun-exposed mosquito wings also reflect UV wavelengths which could be even more attractive than the previously tested white or blue lights (see above). As mosquitoes can sense UV light^39^ and behaviorally respond to it when they seek floral nectar^18^ or oviposition sites,^40^ it is conceivable that UV light reflections play a role in the context of mate recognition.

Volatile or contact sex pheromones have been hypothesized to contribute to mate location and recognition in mosquitos^41—43^ but supportive evidence for such pheromones remains scant. ^26^ Females of *Ae. aegypti* reportedly produce a 3-component sex pheromone blend comprising 2,6,6-trimethylcyclohex-2-ene-1,4-dione (‘ketoisophorone’), 2,2,6-trimethylcyclohexane-1,4-dione (the saturated analogue of ketoisophorone), and 1-(4-ethylphenyl) ethanone (‘ethanone’).^26^ In laboratory but not field settings, ketoisophorone alone elicited swarming-like flight by males. Both ketoisophorone and its saturated analogue prompted “excited flights” by females, whereas ethanone attracted females.^26^

All the visual, acoustic or pheromonal mate location or recognition cues of mosquitoes described above were studied focused invariably on a single sensory modality, discounting possible interactions between cues and their potential function at spatially different scales. Furthermore, specifics of light flash cues on mate attraction such as the number of mosquitoes in a mating swarm generating these cues, or the most attractive wavelengths of these cues, have not yet been experimentally tested. Conceivably, large mating swarms with many mosquitoes ‘flashing lights’ are more attractive than small ones. Conversely, one would predict that mosquitoes swarming at dusk when light flash cues are less conspicuous may not rely on visual cues for mate location or recognition.

Working with diurnal *Ae. aegypti*, we tested six hypotheses (H): (H1, H2) Swarming mosquitoes produce light flashes, and the attractiveness of a mating swarm (i.e., array of light-flashing LEDs) is dependent upon both swarm size (i.e., number of LEDs in array) and the spectral composition of wing flashes (i.e., light emitted by LEDs); (H3) wingbeat light flashes and sound of *Ae. aegypti* females are long- and short-range male attraction signals, respectively; (H4) swarm pheromone of *Ae. aegypti* females increases the attractiveness of their wingbeat light flashes and sound; and (H5) wingbeat light flashes of *Ae. aegypti* males attract mate-seeking females. Working with *C. pipiens* as a model species for nocturnal mosquitoes, we further tested the hypothesis (H6) that dusk-swarming *C. pipiens* do not exploit wingbeat light flashes for mate attraction.

## RESULTS

### H1: Swarming mosquitoes produce light flashes, and the attractiveness of a mating swarm (i.e., array of light-flashing LEDs) is dependent upon swarm size (i.e., number of LEDs in array)

High-speed video recordings of *Ae. aegypti* males swarming in a laboratory setting revealed light flashes reflecting off their wings ([Videos S1](https://data.mendeley.com/datasets/6gvs94p6r2/3/files/c50c07e6-578f-4a95-99d6-49cb87aaae80), [S2](https://data.mendeley.com/datasets/6gvs94p6r2/3/files/b6f88db9-7ddf-4cb5-b939-54160fcfb04e), [S3](https://data.mendeley.com/datasets/6gvs94p6r2/3/files/92b8096f-4609-4ae1-9dda-bb7500c1f5a8), [S6](https://data.mendeley.com/datasets/6gvs94p6r2/3/files/227160ac-1679-4e17-b54f-45c317a3f95f); see Supplementary Material), as also evident by rapid changes of wing-reflected light intensity over time (Fig. 1). *Ae. aegypti* male flight appears to contain a second harmonic of 1854 Hz (Fig. 1), which roughly corresponds to the third harmonic (1995 Hz) of the estimated female wingbeat frequency of 665 Hz. This corroborates previous evidence of the “harmonic convergence” phenomenon.^30,32,35^ The strength of these changes is reduced somewhat by the highly reflective abdomen of *Ae. aegypti*. Unlike in blowflies^37^, the strength of these flashes did depend on viewing angle. As compared with other insects the amplitude of the wing movement forward and backward is small^44^, presenting very little wing surface to reflect light when viewed from the side. While high-speed video recordings of *Ae. aegypti* males swarming in outdoor settings were not obtainable, we documented the ‘wing flash phenomenon’ with other dipterans swarming in a sunlit courtyard ([Videos S4](https://data.mendeley.com/datasets/6gvs94p6r2/3/files/4aa60c23-7bd9-4f06-afc1-be366d4b27d8), [S5](https://data.mendeley.com/datasets/6gvs94p6r2/3/files/58b9680e-e4a8-4ecf-b043-d5b37f6fee9b), Fig. S1). Ablated wings of *Ae. aegypti* males reflected broadly between 300-700 nm, with greater proportional reflection of wavelengths above 500 nm (Fig. S2), likely due to dark brown hairs on the wings.^45^

**Figure 1.**
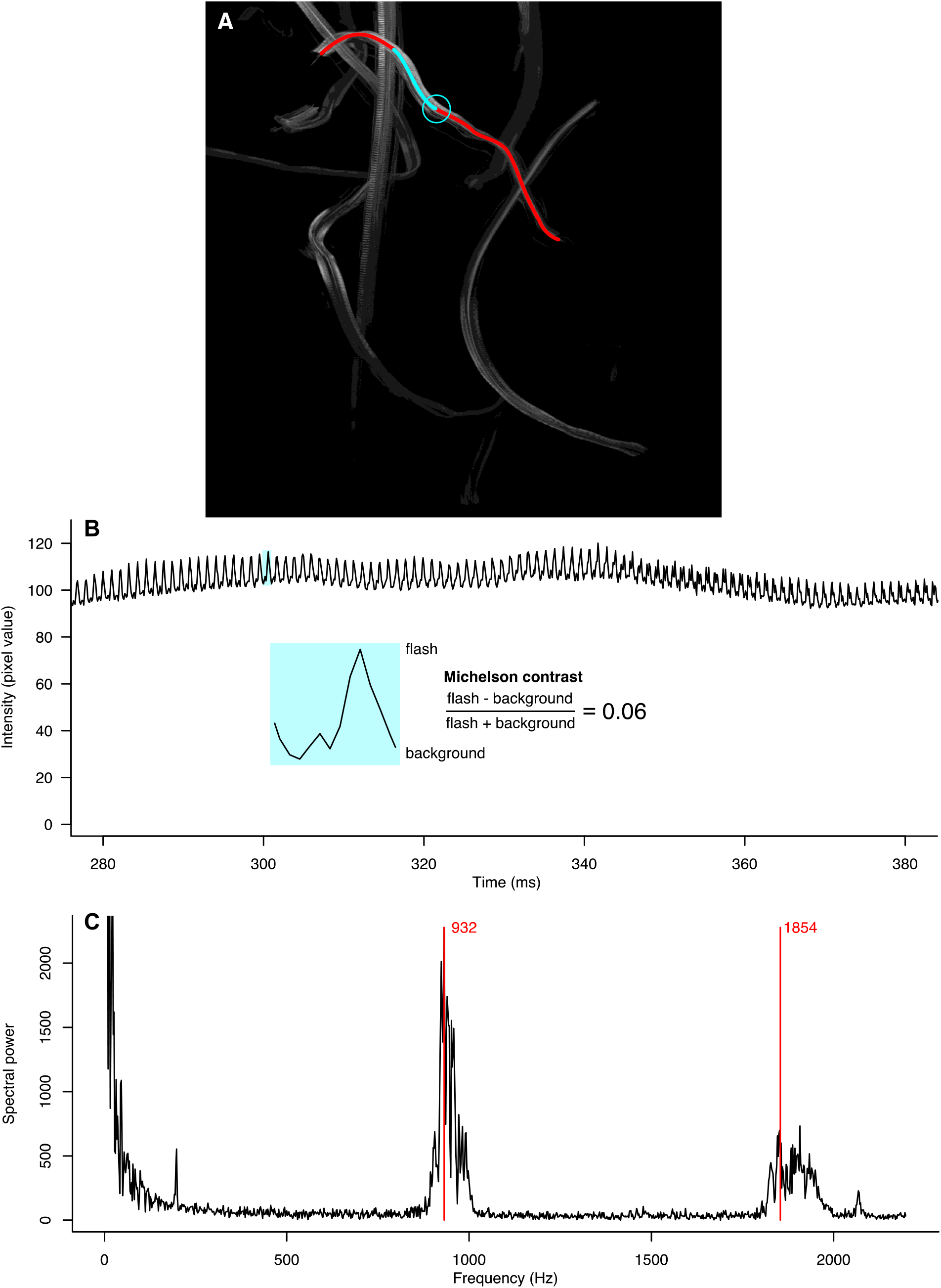
Contrast and frequency analyses of the wing flash series produced by a single male *Aedes aegypti* in a laboratory swarm recorded in Video S1. (A) Z-projection showing the maximum intensity in each pixel over all frames that captured the flight path of the male responding to a 665 Hz tone. The blue track shows the section of the flight path analysed in B, and the red track shows the entire flight path analysed in C. The blue circle indicates the start of the flight path shown in B, and delineates the area (tracking the flying insect) used to characterize the intensity in each frame of this section. (B) Intensity trace showing the mean pixel value within the blue circle across each frame of this section. The two blue insets show a single flash along with a calculated Michelson contrast. (C) Fast Fourier transform periodogram showing the relative spectral power of the wing beat frequency and its 2^nd^ harmonic labeled in red.

To determine the effect of *Ae. aegypti* swarm size on swarm attractiveness, we assembled LEDs in an array, released groups of 50 males into mesh cages for each bioassay, and video recorded their alighting responses (as a measure of attraction) on each of two LED arrays (Fig. 2B) that differed in the number of LEDs flashing blue light (Fig. S8) at 665 Hz (the wing beat frequency of females) (Table 1). When given a choice between a 1-LED array and an 8-LED array, males alighted more often on the latter (F = 67.529, p < 0.0001; Fig. 3, Exp. 1). In contrast, 4- and 8-LED arrays prompted similar numbers of alighting responses by males (F = 0.64, p = 0.64; Fig. 3, Exp. 2). However, 16-LED arrays received three times more alighting responses than 8-LED arrays (F = 22.63, p = 0.001; Fig. 3, Exp. 3). These data in combination support the hypothesis that swarm size affects its attractiveness to mate-seeking males.

**Table 1.** Details of signals [wingbeat light flash, wingbeat sound, pheromone (Phero)] tested in small-space (SS: 61 × 61 × 61 cm) and large space (LS: 2.25 × 2.1 × 2.4 m) behavioral bioassays (see Fig. 2 for experimental design) with *Aedes aegypti* (Exps. 1-11) and *Culex pipiens* (Exps. 12-14)

| Stimulus 1 |  |  |  | Stimulus 2 |  |  |
| --- | --- | --- | --- | --- | --- | --- |
| Exp. #<br>Range | Light flash | Sound | Phero | Light flash | Sound | Phero |
| **H1:** The attractiveness of an *Ae. aegypti* mating swarm (array of light-flashing LEDs) depends upon the number of mosquitoes in the swarm (number of LEDs in the array) |  |  |  |  |  |  |
| 1<br>SS | 1 LED<br>(blue; 665 Hz) | Silent | No | 8 LED<br>(blue; 665 Hz) | Silent | No |
| 2<br>SS | 4 LEDs<br>(blue; 665 Hz) | Silent | No | 8 LEDs<br>(blue; 665 Hz) | Silent | No |
| 3<br>SS | 16 LEDs<br>(blue; 665 Hz) | Silent | No | 8 LEDs<br>(blue; 665 Hz) | Silent | No |
| **H2:** The attractiveness of an *Ae. aegypti* mating swarm (i.e., array of light-flashing LEDs) is dependent upon the spectral composition of wing flashes (i.e., light emitted by LEDs) |  |  |  |  |  |  |
| 4<br>SS | 8 LEDs<br>(UV; 665 Hz) | Silent | No | 8 LEDs<br>(blue; 665 Hz) | Silent | No |
| **H3:** Wing beat light flashes and sound of *Ae. aegypti* females are long- and short-range male attraction signals, respectively |  |  |  |  |  |  |
| 5<br>SS | 8 LEDs<br>(blue; 665 Hz) | Wingbeat sound<br>(female: 665 Hz) | No | 8 LEDs<br>(blue; constant) | Wingbeat sound<br>(female: 665 Hz) | No |
| 6<br>LS | 8 LEDs<br>(blue; 665 Hz) | Wingbeat sound<br>(female: 665 Hz) | No | 8 LEDs<br>(blue; constant) | Wingbeat sound<br>(female: 665 Hz) | No |
| 7<br>SS | 8 LEDs<br>(blue; 665 Hz) | Wingbeat sound<br>(female: 665 Hz) | No | 8 LEDs<br>(blue; 665 Hz) | White noise | No |
| 8<br>SS | 8 LEDs<br>(blue; 665 Hz) | Wingbeat sound<br>(female: 665 Hz) | No | 8 LEDs<br>(blue; 665 Hz) | Silent | No |
| 9<br>SS | 8 LEDs<br>(blue; 715 Hz) | Wingbeat sound<br>(male: 715 Hz) | No | 8 LEDs<br>(blue; 715 Hz) | White noise | No |
| **H4:** Swarm pheromone of *Ae. aegypti* females increases the attractiveness of their wingbeat light flashes and sound |  |  |  |  |  |  |
| 10<br>LS | 8 LEDs<br>(blue; 665 Hz) | Wingbeat sound<br>(female: 665 Hz) | Yes<sup>a</sup> | 8 LEDs<br>(blue; 665 Hz) | Wingbeat sound<br>(female: 665 Hz) | No<sup>b</sup> |
| **H5:** Wing beat light flashes of *Ae. aegypti* males are attractive to mate-seeking females |  |  |  |  |  |  |
| 11<br>SS | 8 LEDs<br>(blue; constant) | No | No | 8 LEDs<br>(blue; 715 Hz) | No | No |
| **H6:** Dusk-swarming *C. pipiens* do not use wingbeat light flashes for mate recognition |  |  |  |  |  |  |
| 12<br>SS | 8 LEDs<br>(white; 350 Hz) | No | No | 8 LEDs<br>(white; constant) | No | No |
| 13<br>SS | 8 LEDs<br>(white; 550 Hz) | No | No | 8 LEDs<br>(white; constant) | No | No |
| 14<sup>c</sup><br>SS | 8 LEDs<br>(white; 550 Hz) | No | No | 8 LEDs<br>(white; constant) | No | No |

<sup>a</sup> ketoisophorone (300 µg) in pentane-ether (30 µl) applied onto filter paper; <sup>b</sup> pentane-ether (30 µl) applied onto filter paper. <sup>c</sup> Exp. 14 was run at low room lighting (1 lux) and with dimmed LEDs ( $6.67 \times 10^{12}$ photons/cm<sup>2</sup>/s).

**Figure 2.**
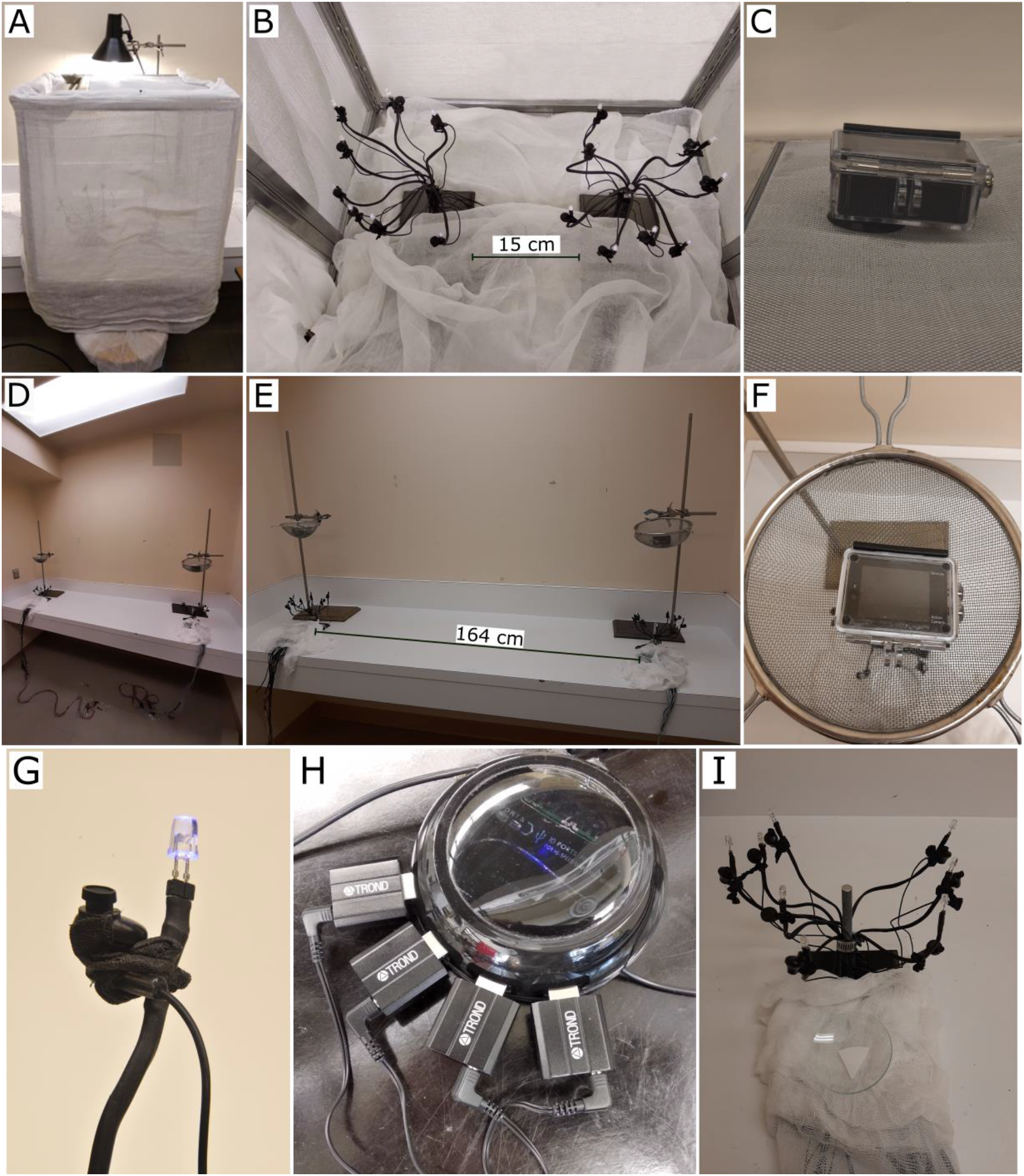
Photographs illustrating the experimental design for testing mosquitoes in behavioural bioassays. (A-C) External and internal views of the small-space bioassay arena (wire mesh cage: 61 × 61 × 61 cm), depicting two assemblies of eight light emitting diodes (LED) each (B), and a video camera on top of the cage (C) for recording alighting responses of mosquitoes on LED assemblies. (D-F) Views of the large-space bioassay room (225 × 210 × 240 cm), with a video camera inside a metal sieve (F) positioned above each of two widely-spaced LED assemblies. The sieve blocked potential electromagnetic waves emanating from the camera. Light was provided via two fluorescent bulbs in the ceiling fixture (for spectral composition see Supplementary Information). (G-I) Details of the experimental design showing a paired LED/earbud speaker mounted on a single arm of the 8-LED assembly (G), the USB hub with USB sound cards driving earbud speakers (H), and a glass dish containing a piece of pheromone- or solvent-treated filter paper (I) deployed in a pheromone experiment.

**Figure 3.**
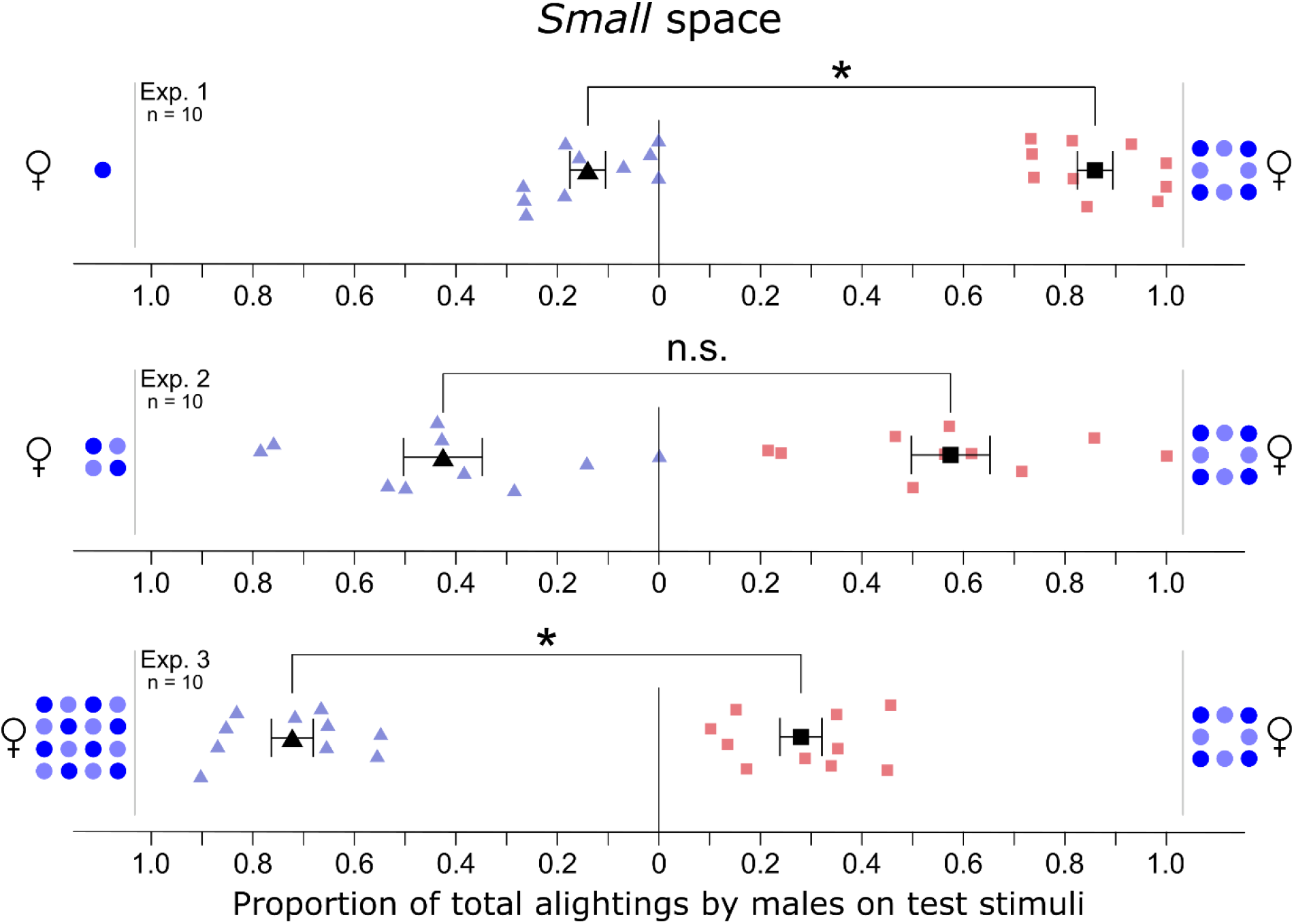
Effect of LED numbers in assemblies on alighting responses of 2- to 7-day-old male *Aedes aegypti*. Numbers of blue dots represent the number of LEDs contained within each of two LED arrays (Fig. 2B) flashing blue light at the 665 Hz wingbeat frequency of female *Ae. aegypti*. Each replicate was run with 50 males. Light blue triangles and light red squares show the data of individual replicates and black symbols the mean (± SE). An asterisk indicates a significant preference (binary logistic regression model; p < 0.05; n. s. = not significant).

### H2: The attractiveness of a mating swarm (i.e., array of light-flashing LEDs) is dependent upon the spectral composition of wing flashes (i.e., light emitted by LEDs)

To determine whether the attractiveness of mating swarms depends upon the wavelength of light reflected off the wings of swarming mosquitoes, we offered groups (n = 10) of 50 males a choice between two 8-LED arrays (Fig. 2B) flashing (665 Hz) either blue light (422 nm) or UV light (360 nm) (Table 1; Fig. S8). Video-recordings revealed that males alighted similarly often on the UV LED array and the blue LED array (n = 10, F = 3.84, p = 0.081; Fig. S3, Exp. 4), suggesting that the short-wave spectral content of wing flashes does not modulate the attractiveness of mating swarms.

### H3: Wingbeat light flashes and sound of Ae. aegypti females are long- and short-range male attraction signals, respectively

To determine whether wingbeat light flashes of females (665 Hz) are long-range male attraction signals, we ran an experiment of identical design in both small and large spatial settings (mesh cage, room; Fig. 2A,B,D,E), offering groups (n = 10) of 50 males a choice between two 8-LED arrays separated by 15 cm (mesh cage) or 164 cm (room) (Table 1). The LEDs of array 1 flashed blue light at 665 Hz, whereas the LEDs of array 2 emitted constant blue light. Each LED in both arrays was coupled with an earbud speaker (Fig. 2G) broadcasting female wingbeat sound (665 Hz). In the cage setting, where males are already near mating swarms (i.e., LED arrays) and can hear the wing beat sound, the type of visual stimulus (flashing or constant light) had no effect on alighting responses by males (n = 10, F = 0.86, p = 0.86; Fig. 4, Exp. 5). Conversely, in the room setting, where males still needed to locate mating swarms (i.e., LED arrays), LED arrays flashing blue light prompted 5.8-times more alighting responses by males than LED arrays emitting constant blue light (n = 10, F = 30.43, p = 0.001; Fig. 4, Exp. 6). The data of both experiments combined support the hypothesis that wingbeat light flashes of females attract males at long-range.

**Figure 4.**
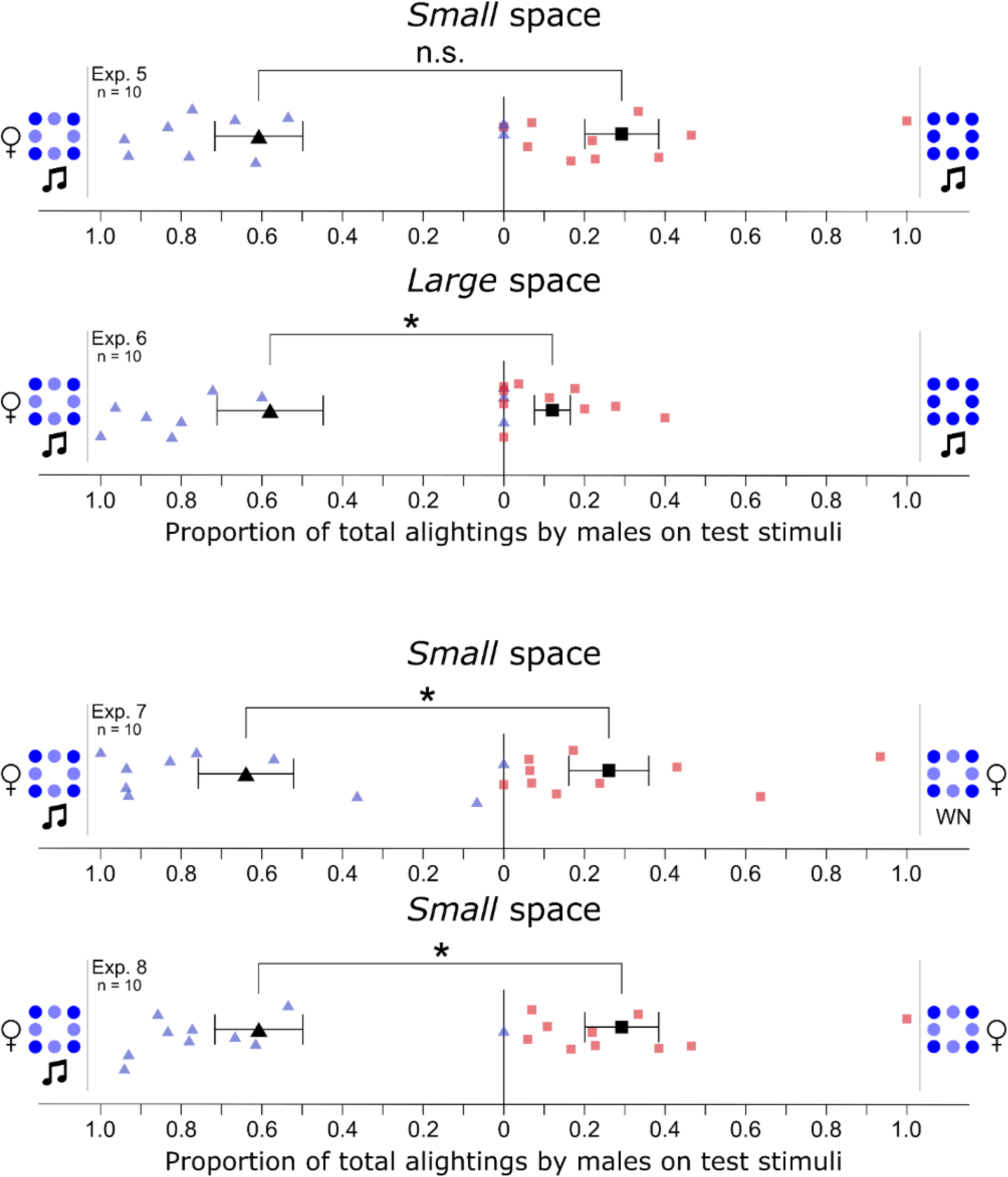
Space-dependent effects of visual and acoustic signals tested in combination on alighting responses of 2- to 7-day-old male *Aedes aegypti*. The number of blue dots represents the number of blue LEDs contained within each of two LED arrays (Fig. 2B), one of which was emitting light flashes (depicted as a mixture of light- and dark-blue dots) at the 665 Hz wingbeat frequency of female *Ae. aegypti*, and the other array was emitting constant light (depicted as uniformly dark-blue dots). Musical notes and WN (white noise) indicate concurrent broadcast of female wingbeat sound (665 Hz) and white noise, respectively. Light blue triangles and light red squares show the data of individual replicates and black symbols the mean (± SE). Experiments were conducted in a mesh cage [*Small* space (Fig. 2A,B); Exps. 5, 7, 8] or within a bioassay room [*Large* space (Fig. 2D,E); Exp. 6]. For each experiment, an asterisk indicates a significant preference (binary logistic regression model; p < 0.05; n. s. = not significant).

To confirm that wingbeat sound of females (665 Hz) is a short-range male attraction signal (see above), we offered groups (n = 10) of 50 males in the mesh cage setting a choice between two 8-LED arrays fitted with earbud speakers that broadcasted either the females’ wingbeat sound (array 1) or white noise (control stimulus; array 2) (Table 1). The LEDs of both arrays flashed blue light (665 Hz). In this cage setting, where males are already near mating swarms (i.e., LED arrays) and can distinguish between arrays with or without wingbeat sound, arrays with wingbeat sound prompted 5-times more alighting responses by males (n = 10; F = 19.87, p = 0.001; Fig. 4; Exp. 7).

To ascertain that the white noise had no repellent effect on the males’ responses in experiment 7, speakers of array 2 were kept silent in follow-up experiment 8 which otherwise was identical (Table 1). Similar to data obtained in experiment 7, arrays with wingbeat sound prompted 4.9-times more alighting responses by males (n = 10, F = 39.97, p = 0.0001; Fig. 4, Exp. 8). The data of both experiments combined support the hypothesis that the wingbeat sound of females attracts males at close range.

To further investigate whether males can indeed distinguish between the wing beat sounds of females and males and are attracted only to the sound of females, we offered groups (n = 10) of 50 males a choice between two 8-LED arrays flashing blue light at 715 Hz (the wing flash frequency of males), with earbud speakers of array 1 emitting male wingbeat sound (715 Hz) and speakers of array 2 broadcasting white noise (Table 1). Fewer alighting responses by males on arrays coupled with male wing beat sound (n = 10, F = 5.49, p = 0.043; Fig. 5, Exp. 9) indicate that males are put off by their own wingbeat sound, obviously distinguishing it from that of females.

**Figure 5.**
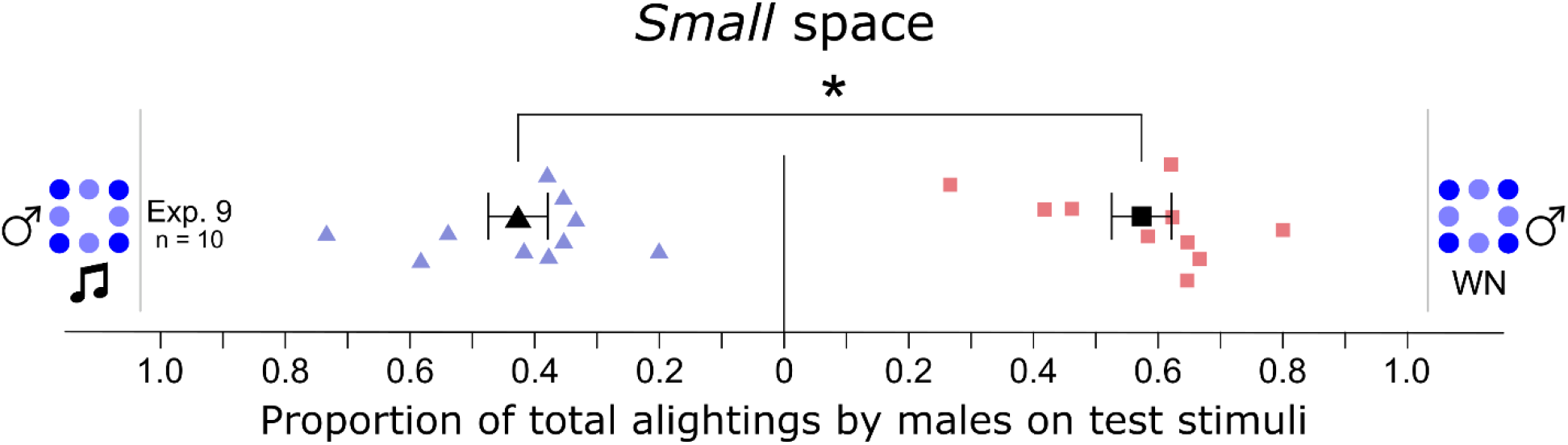
Effect of visual and acoustic signals tested in combination on the alighting responses of 2- to 7-day-old male *Aedes aegypti*. The number of blue dots represents the number of blue LEDs contained within each of two LED arrays (Fig. 2), with LEDs flashing light at the 715 Hz wingbeat frequency of males. The musical note and WN (white noise) indicate concurrent broadcast of male wingbeat sound (715 Hz) and white noise, respectively. Light blue triangles and light red squares show the data of individual replicates and black symbols the mean (± SE). The asterisk indicates a significant preference for WN (binary logistic regression model; p < 0.05).

### H4: Swarm pheromone of Ae. aegypti females increases the attractiveness of their wingbeat light flashes and sound

To test whether the swarm pheromone component ketoisophorone increases the attractiveness of the females’ wingbeat light flash and sound signals, we released groups (n = 9) of 50 males into a room and offered them a choice between two well-spaced 8-LED arrays each fitted with 8 earbud speakers (Fig. 2G, Table 1). All 16 LEDs flashed blue light (665 Hz) and all earbud speakers broadcasted corresponding wingbeat sound (665 Hz). The randomly assigned treatment array was baited with ketoisophorone. Video recording revealed similar numbers of alightings by males on arrays with or without pheromone (n = 9, F = 0.076, p = 0.79; Fig. S4, Exp. 10), indicating no effect of female pheromone on mate-seeking males.

### H5: Wing beat light flashes of Ae. aegypti males are attractive to mate-seeking females

To determine whether wing beat light flashes of *Ae. aegypti* males (715 Hz) attract mate-seeking females, we ran a small-space (cage) experiment, offering groups (n = 13) of 50 females a choice between two 8-LED arrays separated by 15 cm (Table 1). All LEDs of array 1 emitted constant white light (Fig. S8), whereas all LEDs of array 2 flashed white light at 715 Hz. Video-recordings revealed that females alighted more often on arrays with flashing lights than on arrays with constant light (n = 13, F = 4.94, p = 0.046, Fig. 6, Exp. 11).

**Figure 6.**
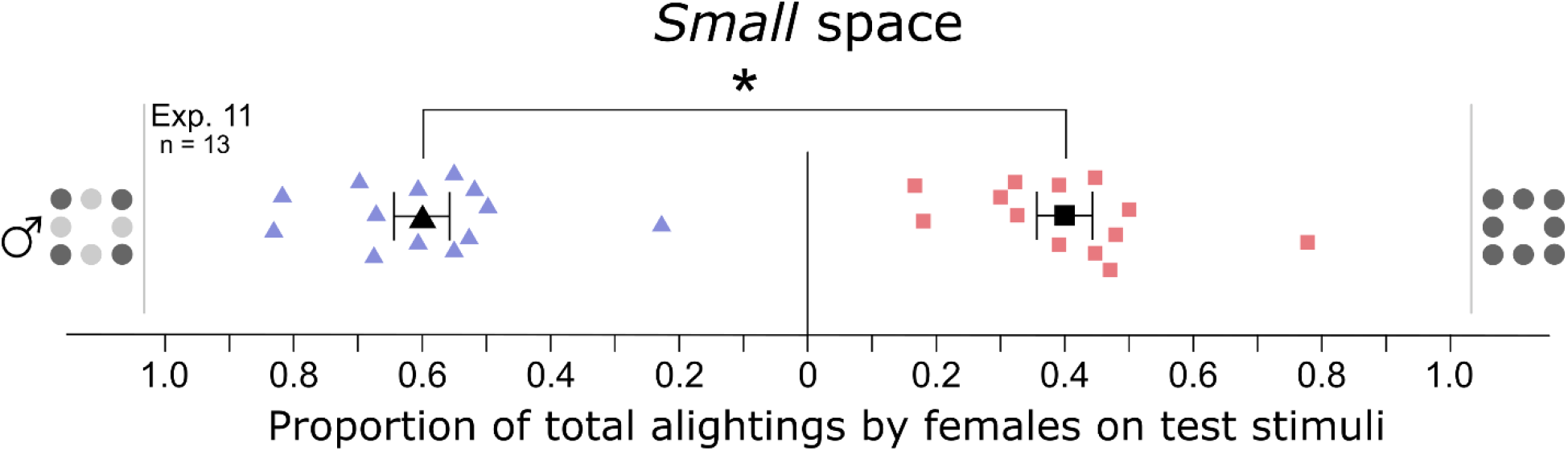
Effect of visual signals on alighting responses of 5- to 10-day-old virgin female *Aedes aegypti*. The numbers of grey dots represent the number of white LEDs contained within each of two LED arrays (Fig. 2), one of which was emitting light flashes (depicted as a mixture of light- and dark-grey dots) at the 715 Hz wingbeat frequency of male *Ae. aegypti*, and the other LED array was emitting constant light (depicted as uniformly dark-grey dots). The asterisk indicates a significant preference for the 715 Hz LEDs (binary logistic regression model; p < 0.05).

### H6: Dusk-swarming C. pipiens do not exploit wingbeat light flashes for mate attraction

To determine whether dusk-swarming *C. pipiens* use wingbeat light flashes as mate attraction signals, we ran three experiments (Exps. 12-14) in the mesh cage setting, one of which (Exp. 14) under dim light (1 lux) (Table 1). In each experiment, we offered groups of 50 2- to 7-day-old *C. pipiens* males a choice between two 8-LED arrays separated by 15 cm. All LEDs of array 1 emitted constant white light (Fig. S8), whereas all LEDs of array 2 flashed white light at either 350 Hz (Exp. 12, n = 10) or 550 Hz (Exp. 13, n = 9; Exp. 14, n = 10), two previously reported wingbeat frequencies of female *C. pipiens*.^34,46^ In all three experiments, very few males alighted on arrays (Fig. S5), revealing no effect of light cues on male attraction, and not warranting statistical analyses of data.

## DISCUSSION

The wing light flash-guided mate location and recognition system of *Ae. aegypti* takes place in a swarm context but otherwise resembles that of other dipterans. This remarkable mate recognition system hinges upon the immense processing speed of dipteran photoreceptors^47,48^ and was only recently discovered in the common green bottle fly, *L. serricata*.^37^ Ever since, the same type of system has been shown to occur in other dipteran taxa, including house flies, *Musca domestica*, black soldier flies, *Hermetia illucens*, and *Ae. aegypti*.^27^

The system in green bottle flies depends upon both the frequencies of light flashes caused by moving wings being sex- and age-specific, and the ability of male bottle flies to recognize the light flash frequency of young female flies that are prospective mates.^37^ A single LED flashing white light at the wingbeat frequency of young females (178 Hz) is sufficient to attract and prompt alighting responses by males.^37^ In *Ae. aegypti*, however, mate location typically takes place in a swarm context,^22,49^ and a single light-flashing LED is not attractive to males or females (Gries et al., unpubl.). To present a ‘mating swarm’ and to test its attractiveness to males, we built assemblies of 8 LEDs (Fig. 2) and offered groups of males a choice between two assemblies that emitted either constant light or light flashing at one of eight frequencies (430, 480, 500, 545, 665, 800, 950 Hz).^27^ In these experiments, males invariably alighted more often on flashing-light LEDs than on constant-light LEDs (Fig. S6; adapted from^27^), suggesting that mate-seeking males may respond to flashing lights of swarming males to locate swarms. However, the effect of wingflash light signals on the responses of males in this previous study^27^ was tested in the absence of wingbeat sound and in a relatively small space. To reveal the effects of light and sound signals [which are perceived at long and short (< 25 cm) range, respectively] at different spatial levels, we ran experiments in both a large setting (2.25 × 2.1 × 2.4 m high) and a small setting (61 × 61 × 61 cm). Our selection of the female (rather than the male) wingbeat light flash and sound frequency (665 Hz each) as test stimuli for the response of males was guided by four considerations: (1) even though females do not form mating swarms on their own, multiple females may concurrently be present in a mating swarm sought after by males. For example, in *Anopheles stephensi mysorensis*, as many as 23% of swarm mates were found to be females;^50^ (2) males ought to be able to recognize females approaching a swarm, or flying well apart within a swarm, at a distance greater than the hearing range for wingbeat sound (15-25 cm);^36^ (3) light flash frequencies covering the range produced by females (665 Hz) and males (715 Hz) were both highly and almost equally attractive to males (Fig. S6); and (4) mate location in *Ae. aegypti* may also occur in a context other than mating swarms.^42,51^

Our data show that flashing lights (665 Hz) are long-range signals that attract males to mating swarms or to mates (Fig. 4). In a large-space setting, LED assemblies flashing light at 665 Hz and emitting wingbeat sound (665 Hz) prompted 5.8-times more alighting responses than LED assemblies emitting constant light and wingbeat sound (665 Hz) (Fig. 4, Exp. 6). Conversely, in a small space setting, when wingbeat sounds were present, flashing lights had no apparent signal characteristics. Each of two LED assemblies producing either flashing or constant light induced similar numbers of alightings by males (Fig. 4, Exp. 5).

Our data (Fig. 4, Exps. 7, 8) also confirm that the wingbeat sound of females is a close-range signal to mate-seeking males.^36,52,53^ When offered a choice between two LED assemblies, both flashing light (665 Hz) but only one emitting female wingbeat sound, males alighted 5.0- and 4.9-times more often on assemblies emitting female wingbeat sound than on assemblies that emitted white noise or were silent (Fig. 4, Exps. 7, 8). Conversely, the wingbeat sound of males (715 Hz) was off-putting to mate-seeking males (Fig. 5, Exp. 9), corroborating previous conclusions that males distinguish between wingbeat sounds of females and males.^25,54^

The attractiveness of light flash mate location signals – tested in small-space bioassays in the absence of sound signals – is modulated not only by the flash frequency (Fig. S6) but also by the number of signals (i.e., mosquitoes in mating swarms, or LEDs in assembly) and the wavelengths of flashing lights. Increasing the number of LEDs in assemblies increased the number of mosquitoes alighting on assemblies (Fig. 3, Exps. 1-3), suggesting that larger mating swarms, or swarms containing a higher percentage of females, are more attractive to mate-seeking males. LED assemblies emitting UV light were as attractive to males as blue-light LED assemblies (Fig. S3, Exp. 4) which were more attractive than white-light LED (Fig. S7) assemblies. Whether equivalent physical characteristics of visual mate location signals affect the behaviour of females is not yet known. However, our findings that females, on average, alighted more often on LED assemblies flashing light at the male wingbeat frequency (715 Hz) than on LED assemblies emitting constant light (Fig. 6, Exp. 11), suggest that females may recognize a mating swarm, in part, based on the flashing lights ‘produced’ by swarming males.

With convincing data showing that visual and acoustic signals contribute to long- and short-range mate location in *Ae. aegypti* (Fig. 4), there was ample incentive to also test the effect of a chemical signal, the female-produced pheromone,^26^ on responses of males. We predicted that female pheromone presented in combination with light and sound signals would modulate the behavior of males. However, the synthetic pheromone component ketoisophorone added to the bimodal complex of visual sound signals failed to express any additive or synergistic effect on the responses of males (Fig. S4, Exp. 10). It is conceivable, though, that the still-air setting of this experiment, with pheromone dissemination being entirely reliant on diffusion without forming a discrete pheromone plume, was not conducive for male attraction. Alternatively, in the absence of air current, pheromone may have built up in the room, ultimately disorienting males rather than guiding them to the pheromone source.

The sensory drive theory predicts functional links between signal design and presentation such that the conspicuousness of signals is maximized relative to environmental conditions and background noise.^55^ Previous reports in the literature and our data on *Ae. aegypti* and *C. pipiens* are in complete agreement with these predictions. The onset of the photophase induces swarm formation by male *Ae. aegypti*.^43^ With incident light reflecting off the wings of swarming males, their swarm becomes a visual beacon for other males and females in search for mates. As more mosquitoes enter the swarm, the “firework” of light flashes becomes larger and more attractive (Fig. 3; Videos S1–S6). The conspicuousness of the swarm display is further enhanced 3- to 4-times when putting the light flash LED assembly on an oscillating shaker table,^27^ mimicking a swarm gently swaying in the wind. In contrast, visual mate location systems hinging on incident sunlight reflecting off the wings of in-flight dipterans, as shown for bottle flies, house flies and black soldier flies,^27,37^ as well as yellow fever mosquitoes^27^ (also shown in this study), would not be expected to evolve in crepuscular mosquito species such as *C. pipiens* that swarm at dusk when sunlight is absent and illumination is dominated by diffuse light from the horizon.^56^ As predicted, LED assemblies flashing light at the reported wingbeat frequencies of *C. pipiens* (350 Hz,^46^ 550 Hz^34^) had no signal characteristics for bioassay mosquitoes and prompted hardly any behavioral responses (Fig. S5; Exps. 12–14).

In conclusion, we describe that mate location or recognition in *Ae. aegypti* is mediated, in part, by long-range wingbeat light flash signals and by short-range wingbeat sound signals. The attractiveness of the light flash signals is dependent upon both the number of light flashes (i.e., mosquitoes in the swarm) and the wavelengths of the flashing light (i.e., light reflected off wings). As both male and female *Ae. aegypti* respond to light flash signals, these signals apparently contribute to the processes of forming and locating mating swarms. Moreover, with males and females having significantly different wingbeat frequencies,^32,57^ and thus light flash frequencies, the flash frequency could also facilitate long-range recognition of prospective mates. Our data address knowledge gaps as to how male and female *Ae. aegypti*, and possibly the sexes of other (diurnal) mosquitoes, find each other.^31^ Elucidating the mate location and courtship biology of mosquitoes will inform quality assessments of males that are mass-reared and released in sterile insect release tactics. Successful integration of these tactics into mosquito vector control programs^58—60^ hinges on sterile and transgenic males effectively competing with wild males for access to females.

## Supporting information

Supplemental materials for main text

## ACKNOWLEDGEMENTS

We thank Pawel Kowalski and Anthony Slater (electronics shop, SFU), and James Shoults (machine shop, SFU), for assistance in designing and building experimental equipment, Amanda Brooks for contributions to mosquito rearing and data collection, Regine Gries (RG) for preparing pheromone lures, GG and RG for feeding mosquitoes on their arms, and Randeep Sood (High Speed Imaging Inc.) for technical assistance with the high-speed camera.

## AUTHOR CONTRIBUTIONS

EK and GG conceived the study; EK and CL ran behavioural bioassays and scored data; AB and EK obtained high-speed video recordings; AB and EK analyzed data; AB, EK and CL obtained spectrometric data of visual test stimuli; ST and EK produced and adjusted audio files; EK sourced and set up bioassay equipment; EK and GG wrote the first draft, and all authors reviewed and approved of the final draft.

## METHODS

### KEY RESOURCES TABLE

| REAGENT or RESOURCE | SOURCE | IDENTIFIER |
| --- | --- | --- |
| Experimental models: organisms/strains |  |  |
| *Culex pipiens* laboratory colony | This paper | N/A |
| *Aedes aegypti* laboratory colony | This paper | N/A |
| Software and algorithms |  |  |
| Audacity | Audacity Team<sup>61</sup> | https://www.audacityteam.org/ |
| FIJI | Eliceiri/LOCI group<sup>62</sup> | https://imagej.net/software/fiji/ |
| mpv | mpv<sup>63</sup> | https://mpv.io/ |
| R | R Core Team<sup>64</sup> | https://www.R-project.org/ |
| Raspbian 10 | Raspberry Pi Foundation, Cambridge, UK | https://www.raspberrypi.org/software/operating-systems/ |
| SpectraSuite | Ocean Optics | https://www.oceaninsight.com/support/software-downloads/ |
| TrackMate | https://imagej.net/plugins/trackmate/<sup>65</sup> | https://github.com/fiji/TrackMate |
| Other |  |  |
| R code for statistical analysis | Associated Mendeley Data dataset<sup>66</sup> | https://doi.org/10.17632/6gvs94p6r2.3 |
| 665 Hz sounds files used in acoustic experiments | Associated Mendeley Data dataset<sup>66</sup> | https://doi.org/10.17632/6gvs94p6r2.3 |
| 715 Hz sound files used in acoustic experiments | Associated Mendeley Data dataset<sup>66</sup> | https://doi.org/10.17632/6gvs94p6r2.3 |
| Raspberry Pi 3b+ | Cana Kit Corporation | https://www.canakit.com/raspberry-pi-3-model-b-plus-starter-kit.html |
| 5218268F White LEDs | Dialight | http://www.dialightsignalsandcomponents.com/Product/Details/445142/474752 |
| TLHB5800 Blue LEDs | Vishay Intertechnology | https://www.vishay.com/docs/83021/tlhb5800.pdf |
| EOLD-355-525 UV LEDs | OSA Opto Light GmbH | https://www.osa-opto.com/smd-leds/eold-355-525.html |
| HPX-2000 xenon light source | Ocean Optics | https://www.oceaninsight.com/support/documents-manuals/manuals-operating-instrs/hpx-2000-manual/ |
| HR-4000 Spectrophotometer | Ocean Optics | https://www.oceaninsight.com/products/spectrometers/high-resolution/hr-series/ |
| Gould 20Ms/sec Digital Recording Oscilloscope | Gould Electronics GmbH | N/A |
| Photron FASTCAM NOVA S16 | Photron USA, Inc. | https://photron.com/fastcam-nova-s/ |

### Lead Contact

Correspondence and requests for materials should be sent to the Lead Contact, Elton Ko

### Materials Availability

This study did not generate new unique reagents.

### Data and Code Availability

Raw data and code are available on Mendeley Data^66^: https://doi.org/10.17632/6gvs94p6r2.3

## EXPERIMENTAL MODEL AND SUBJECT DETAILS

### Rearing of experimental insects

*Aedes aegypti* mosquitoes were reared in the insectary of the Burnaby campus of Simon Fraser University (SFU) at 23–28 °C, 40-60% RH, and a photoperiod of 14L:10D. Adult mosquitoes were kept in mesh cages (30 × 30 × 46 cm high) provisioned with a 10-% sucrose solution *ad libitum* and allowed to blood-feed on the arm of GG or Regine Gries once a week. Three days after blood-feeding, gravid females were offered an oviposition site consisting of a 354-mL water-filled paper cup (Solo Cup Company, Lake Forest, IL, USA) lined with a paper towel (Kruger Inc., Montreal, QC, Canada). For storage, egg-lined towels were inserted into Ziploc bags (S.C. Johnson & Son, Inc., Racine, WI, USA) kept at 23–28 °C. To initiate a new generation of mosquitoes, towels were transferred to a glass dish (10 cm diam × 5 cm high), containing water enriched with brewer’s yeast (U.S. Biological Life Sciences, Salem, MA, USA). After egg hatching, 1^st^ instar larvae were transferred to water-filled trays (45 × 25 × 7 cm high) and provisioned with NutraFin Basix tropical fish food (Rolf C Hagen Inc., Montreal, QC, Ca). Using a 7-mL plastic pipette (VWR International, Radnor, PA, USA), pupae were transferred to water-filled Solo Cups covered with a mesh lid and fitted with a sucrose solution-soaked cotton ball to sustain adult mosquitoes eclosing over the course of 72 h. These mosquitoes were then released into mesh cages (30 × 30 × 46 cm high) and separated by sex for use in bioassays when they were 2–7 days old (males) or 5–10 days old (females).

The rearing protocol for *C. pipiens* resembled that for *Ae. aegypti* except that *(i)* rooms were kept at 23–26 °C, (*ii*) gravid females were offered a glass dish (10 cm diam × 5 cm high) as oviposition site, and (*iii*) egg rafts – rather than egg-lined towels – were transferred to water-filled trays for larval development. Only 2- to 7-day-old males were tested in bioassays.

## METHOD DETAILS

### High-speed videography

Swarming of *Ae. aegypti* males and of taxonomically unidentified (dipteran) midges was video-recorded using a Photron FASTCAM NOVA S16 high-speed camera (Photron USA Inc., San Diego, CA 92126, USA) fitted with a Nikon NIKKOR AF-S Micro lens (105 mm, f/2.8 AF) for close-up shots, or a Nikon NIKKOR Telephoto Zoom lens (AF 35–80 mm, f/4-5.6D) for wide angle shots (both lenses: Nikon Canada Inc., Mississauga, ON, CA). For further magnification, a clip-on macro filter (DCR-250 Super MacroScan Conversion lens, Yoshida Industry Co., Ltd, Tokyo, Japan) was attached to the 105-mm micro lens. The high-speed camera was connected via an ethernet cable to an ASUS laptop computer (ASUS Canada, Markham, ON, CA) running Photron FASTCAM Viewer 4 (PFV4) (Photron USA, Inc., San Diego, CA 92126, USA). Videos were recorded at 3,000, 5,000 or 10,000 frames per second (fps), with shutter speeds of 1/6,000, 1/10,000 or 1/30,000 s. Videos were downloaded from the camera as mRAW files, and converted with PFV4 to MP4 files.

To film swarming behavior of *Ae. aegypti* males, 100 males were released into a mesh cage (12 × 12 × 12 cm) (BioQuip Products Inc., Rancho Dominguez, CA 90220, USA), and exposed to host cues (CO_2_ released from dry ice; EK’s hand or forearm) and mate recognition signals (female wingbeat sound [665 Hz] played back from a smartphone [Video S6]). The cage was illuminated from above with an LED light source (AOS Offboard LED light, AOS Technologies AG, Baden-Daettwil, CH) mounted on a friction arm (Vitec Imaging Solutions Spa, Cassola, IT). One mesh wall of the cage was replaced with plastic wrap to facilitate filming.

Flies swarming in a sunlit courtyard on the Burnaby campus of SFU were video recorded at 16:00 on 01 September from an open, second-story window overlooking the courtyard. Because the midges were swarming too high above ground and too far away from the window, voucher specimens could not be obtained for identification and deposition in a museum collection.

To quantify the light intensity of the insects across their flight paths, frames of the video were imported into FIJI^62^ as 12-bit Tiff images in a manner that preserved sensor linearity.^66^ We used the plugin TrackMate^65^ to follow the position of the insect over their flight path with a circular region of interest with a diameter encompassing both the wingspan and body length of the insect. The mean pixel value was recorded from within this circular region of interest for each frame of the flight path. These intensity data were then imported into R 3.6.2^64^ to calculate fast Fourier transformation periodograms.

### Spectroscopy of wing reflections

Wings of *Ae. aeygpti* males were ablated from live insects and mounted on an insect pin. These wings were illuminated by a xenon light source (HPX-2000, Ocean Optics, Dunedin, FL, USA) via a fibre optic cable, and their reflections were sampled with a cosine corrector connected to a spectrophotometer (HR-4000, Ocean Optics) fitted with a fibre cable. To capture both the minimal and the maximal possible reflection, wings were positioned either edge on with respect to the cosine corrector, or at an angle where their specular reflection was directed at the center of the cosine corrector. For comparison with the wing reflections, the spectra of the xenon light source was also measured using a square of aluminum foil positioned in a similar way to the specularly reflecting mosquito wing.

### LEDs

Spectra of white LEDs (5218268F, Dialight, London, UK), blue LEDs (TLHB5800, Vishay Intertechnology, Malvern, PA, USA) and UV LEDs (EOLD-355-525, OSA Opto Light GmbH, Berlin, DE) (Fig. S8) were recorded with a spectrophotometer (HR-4000, Ocean Optics) and SpectraSuite software (Ocean Optics). The photon flux of each LED was sampled at a distance of 5 cm from the cosine corrector connected to the sampling fibre of the spectrometer. This allowed use to vary the amperage supplied to the LEDs in order to achieve an intensity of 2e^15^ photons/cm^2^/s. Using a lathe, the lens of each LED was flattened to widen the angle of emitted light. The frequency (Hz) and the duty cycle (set to 3%) of each LED were verified using an oscilloscope (Gould 20Ms/sec Digital Recording Oscilloscope, Gould Electronics GmbH, Eichstetten am Kaiserstuhl, DE).

### Design of LED arrays

LED arrays consisted of up to 16 LEDs arranged in a three-dimensional circular shape (∼15 cm diam) (Fig. 2B). Each LED was mounted upward-facing 18–23 cm above ground on a separate, rigid stalk which was attached to a ring stand, the base of which was covered with Cheesecloth (Cheesecloth Wipes, VWR International, PA, USA) to minimize visibility (Fig. 2B). Each LED was connected to one channel of a 16-channel pulse generator (5-Volt, 2-Amp) designed and built by the Science Technical Centre at SFU to allow independent control of test variables for each LED, including duty cycle, frequency (Hz), amperage and periodicity.

### General design of small-space behavioural experiments

We ran behavioural bioassays with mosquitoes in a mesh cage (61 × 61 × 61 cm) (BioQuip Products, Inc., CA, USA) (Fig. 2A,B), with the cage bottom and the front and side walls covered with cheesecloth to minimize stray light entry and light reflectance. A lamp fitted with an LED bulb (Feit Electric, Pico Rivera, CA, USA; Fig. S9) was placed above the rear edge of the cage to provide illumination during bioassays. For each 20-min bioassay, we placed two LED arrays (see above) 15 cm apart from each other in the centre of the cage (alternating their position between replicates) and released 50 2- to 7-day-old sexually mature males (Exps. 1–10, 12–14), or 50 5- to 10-day-old sexually mature females (Exp. 11), into the cage. To video-record alighting responses of mosquitos on LEDs, we placed an AKASO EK7000 action camera (AKASO, Frederick, MD, USA) on top of the cage (Fig. 2C). Mosquito contacts and landings on LEDs, or on a stalk within 2.5 cm of an LED, were recorded as responses. During bioassays, rooms were maintained at a temperature of 23–28 °C and 40-60% relative humidity. After each bioassay, the camera was stopped and the cage was opened to release the mosquitoes which were then euthanized with an electric fly swatter (Guangzhou Sidianjin Trading Co., Guangzhou, CN). To optimize the responsiveness of *Ae. aegypti* females and males in all bioassays (see also below), we tested them only on sunny or overcast (but not rainy) days and only during the light phase of their photoperiod (14L:10D).

### General design of large-space behavioural experiments

In a cubicle (2.25 × 2.1 × 2.4 m high; Fig. 2D) of the insectary illuminated by ceiling fluorescent lighting (F32T8/SPX50/ECO, General Electric, Boston, MA, USA; Fig. S9), two LED arrays were placed on a counter 164 cm apart from each other, and 43 cm and 30 cm, respectively, away from the back and side walls of the cubicle (Fig. 2E). For each bioassay, 50 2- to 7-day-old sexually mature males were released into the cubicle through the cubicle door. Their alighting responses on LEDs were video recorded with an AKASO action camera placed in a metal sieve (shielding the camera’s electromagnetic field) (Fig. 2F) mounted on a ring stand 42 cm above each LED array. During bioassays, rooms were kept at 23-28 °C and 40-60% relative humidity. After 20 min of recordings, the cameras were turned off, all mosquitoes were euthanized with an electric fly swatter, and the position of LED array 1 and 2 was reversed for the next replicate.

### Wingbeat sound cues

To determine the effect of mosquito wing beat sound on LED-alighting responses of bioassay mosquitoes, we used Audacity 2.3.2^61^ to prepare eight sound files^66^ with paired channels, one of which was randomly assigned to the treatment stimulus and the other to the control stimulus. Treatment stimuli consisted of wingbeat sound characteristic of *Ae. aegypti* females (665 Hz) or males (715 Hz), whereas control stimuli consisted of white noise (sound that covers the entire range of audible frequencies). Audio tracks of wing beat frequencies or white noise were played in parallel, Doppler-shifting upwards, holding steady, or Doppler-shifting downwards to silence, each of these three phases lasting 7 s. The intensity level of the wing beat sound and the white noise control stimulus were each adjusted to 10 dBL above background (SPL = 45 dBL), measured 2.5 cm away from each sound-emitting earbud speaker (RPHJE120K, Panasonic, Osaka Prefecture, Japan), using a 1551-C sound level meter fitted with a Type 1560-PB microphone (General Radio Company, Concord, MA, USA). Earbud-emitted sound was not audible to human hearing at 50 cm away from the source. Each headphone pair played back either an artificial tone (665 Hz, 715 Hz), white noise or was kept silent, depending on the array (treatment or control) and the experiment. Sound files were played using MPV media player.^63^

To reduce the directionality of sound stimuli, we removed the rubber tip from each earbud. On both arrays, each of eight LEDs was paired with a single upward-facing earbud which was attached with a twist tie to the LED-carrying stalk 2 cm below the LED (Fig. 2G). Earbud wires on the cage floor were covered with cheesecloth and routed out of the cage through a mesh sleeve. Each pair of earbuds (one earbud being assigned to the treatment array and the other to the control array) was plugged into a separate USB sound card (C-Media HS-100B Chipset, TROND, Shenzhen, CN) which, in turn, was plugged into a 4-port USB hub (Qicent, Shenzhen, CN) (Fig. 2H). Connecting only two soundcards to each of four USB hubs helped avoid latency of playback recordings. The USB hubs were plugged into a Raspberry Pi 3 B+ computer (Cana Kit Corporation, North Vancouver, BC, Ca), running Raspbian 10 (Raspberry Pi Foundation, Cambridge, UK).

### Specific experiments

#### H1: The attractiveness of a mating swarm (i.e., array of light-flashing LEDs) is dependent upon swarm size (i.e., number of LEDs in array)

To determine the effect of LED numbers in array (i.e., ‘mosquito swarm size’) on array attractiveness (Exps. 1–3; n = 10 each; Table 1), we presented groups of 50 *Ae. aegypti* males each with a choice of two LED arrays that differed in number of LEDs. Specifically, we tested arrays with eight *vs* one LED (Exp. 1), eight *vs* four LEDs (Exp. 2), and eight *vs* 16 LEDs (Exp. 3). Each LED in each array flashed blue light at 665 Hz.

#### H2: The attractiveness of a mating swarm (i.e., array of light-flashing LEDs) is dependent upon the spectral composition of wing flashes (i.e., light emitted by LEDs)

The effect of LED wavelength (UV or blue) on LED-alighting responses by *Ae. aegypti* males was tested by offering groups of 50 males each a choice between two 8-LED arrays flashing either UV or blue light at 665 Hz (the light flash frequency of flying females) (Exp. 4, n = 10; Table 1). The amperage supplied to LEDs was modulated to an equal photon flux from the blue and UV LEDs.

#### H3: Wingbeat light flashes and sound of Ae. aegypti females are long- and short-range male attraction signals, respectively

To test whether wingbeat light flashes (665 Hz) of *Ae. aegypti* females are long-range male attraction signals, we ran a two-choice experiment in both a small setting (61 × 61 × 61 cm; Exp. 5, n = 10; Fig. 2A,B) and a large setting (225 × 210 × 240 cm high; Exp. 6, n = 10; Fig. 2D,E; Table1). In each experiment, we offered groups of 50 males each a choice between two 8-LED arrays which were separated by 15 cm (Exp. 5) or 164 cm (Exp. 6). In both experiments, the LEDs of array 1 emitted blue light flashes of 665 Hz, whereas the LEDs of array 2 emitted constant blue light. Each LED in both arrays was coupled with an earbud speaker emitting the females’ wingbeat sound (665 Hz).

To test whether wingbeat sounds (665 Hz) of *Ae. aegypti* females are short-range mate recognition signals for males, we ran a small setting experiment, offering males a choice between two 8-LED arrays separated by 15 cm. The LEDs of both arrays emitted blue light flashes at 665 Hz. The earbud speakers of array 1 emitted female wingbeat sound (665 Hz), whereas speakers of array 2 emitted white noise (Table 1; Exp. 7, n = 10). To determine whether white noise may have had a repellent effect on the males’ responses in Experiment 7, speakers of array 2 were kept silent in follow-up experiment 8 (n = 10) which otherwise was identical (Table 1). To further investigate whether mate recognition cues of males deter males, we offered groups of 50 males each a choice between two 8-LED arrays emitting blue light flashes at 715 Hz (the wing flash frequency of males), with earbud speakers of array 1 emitting male wing beat sound (715 Hz) and speakers of array 2 broadcasting white noise (Table 1; Exp. 9, n = 10).

#### H4: Swarm pheromone of Ae. aegypti females increases the attractiveness of their wingbeat light flashes and sound

To test whether the swarm pheromone component ketoisophorone increases the attractiveness of the females’ wingbeat light flashes and sound, we ran a large setting (room) experiment (Fig. 2D,E; Table 1; Exp. 10, n = 9), offering groups of 50 males each a choice between two 8-LED arrays separated by 164 cm. All LEDs and earbud speakers of both arrays emitted blue light flashes (665 Hz) and the corresponding wingbeat sound (665 Hz). The bases of both arrays were fitted with a filter paper-lined watch glass which was treated with either ketoisophorone (300 μg) in pentane-ether (30 μl) (array 1) or a pentane-ether control (30 μl) (Fig. 2I). The solvent was allowed to evaporate completely prior to the onset of each bioassay.

#### H5: Wing beat light flashes of Ae. aegypti males are attractive to mate-seeking females

To determine whether wing beat light flashes of *Ae. aegypti* males are attractive to mate-seeking females, we ran a small-setting (cage) experiment, offering groups of 50 females each a choice between two 8-LED arrays separated by 15 cm and deprived of all earbud speakers. All LEDs of array 1 emitted constant white light, whereas all LEDs of array 2 emitted white light flashes (715 Hz) (Table 1; Exp. 11, n = 13).

#### H6: Dusk-swarming C. pipiens do not exploit wingbeat light flashes for mate attraction

To determine whether dusk-swarming *C. pipiens* use wingbeat light flashes as mate recognition cues, we ran three small-setting experiments, offering groups of 50 2- to 7-day-old males a choice between two 8-LED arrays separated by 15 cm and deprived of all earbud speakers. All LEDs of array 1 emitted constant white light, whereas all LEDs of array 2 emitted white light flashes at either 350 Hz (Exp. 12, n = 10) or 550 Hz (Exps. 13, 14, n = 10 each; Table 1), two previously reported wingbeat frequencies of female *C. pipiens*.^34,46^ Experiments 12 and 13 followed the ‘general design of small-space behavioural experiments’ (see above).

Taking into account that *C. pipiens* forms mating swarms at dusk, the room lights in follow-up experiment 14 were turned off and the bioassay cage was illuminated from behind by an LED bulb (Feit Electric, Pico Rivera, CA, USA; Fig. S9) set by a dimmer (TBL03, Leviton Manufacturing Company, Inc., Melville, NY, USA) to a light intensity level of 1 Lux at the cage centre.^34^ Likewise, the photon flux of LEDs in arrays 1 and 2 emitting constant light and flashing light, respectively, was reduced to 6.67e^12^ photons/cm^2^/s in accordance with the low light level in the room. To facilitate recordings of alighting responses by mosquitoes on LEDs, we used a hunting camera (Campark Trail Camera, Campark Electronics Co., Ltd, HK) with an IR-sensitive wavelength range which mosquitoes cannot perceive.

### Statistical analyses

We used R 3.6.2^64^ to analyse behavioural data. Mean proportions of contact and alighting responses by mosquitoes were analyzed with logistic regression using generalized linear models. ^66^ In order to determine whether proportions differed between arrays, we compared an intercept only model to a null model with a likelihood ratio test. We then used back-transformed coefficients from those models to obtain mean and standard errors for the proportion of mosquitos responding to each array.

