## Supplemental materials for main text for "Long- and short-range bimodal signals mediate mate location and recognition in yellow fever mosquitoes"

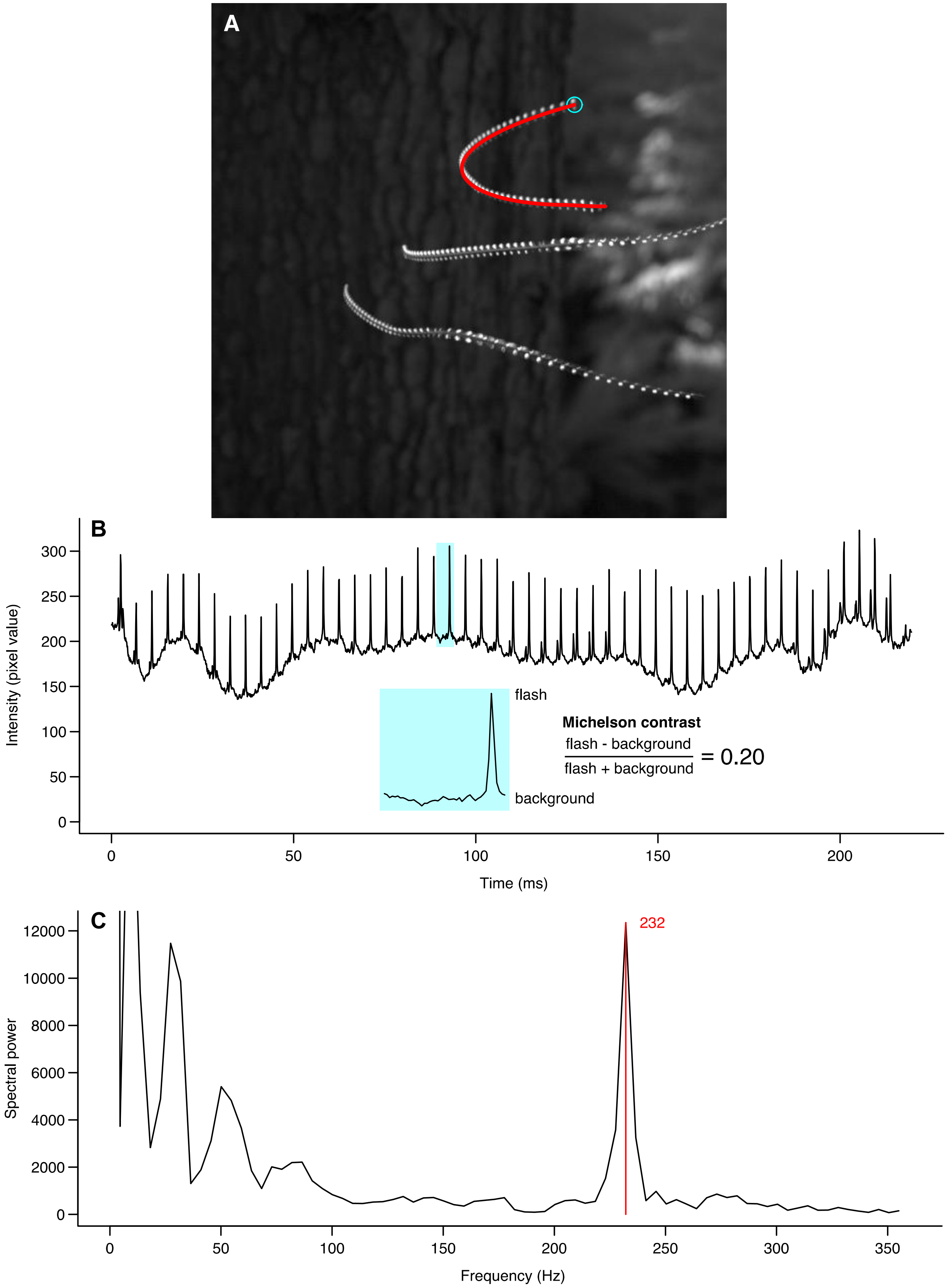
A
B
Intensity (pixel value)
Time (ms)
flash
background
Michelson contrast
flash - background
flash + background
= 0.20
C
Spectral power
Frequency (Hz)
232

**Figure S1. Contrast and frequency analysis of the wing flash series produced by a single insect in an outdoor swarm of midges recorded in Video S4**. (A) Z-projection showing the maximum intensity in each pixel over all frames that captured the insect's flight path. The blue circle indicates the start of the flight path and delineates the area (tracking the flying insect) used to characterize the pixel intensity in each frame. The red track shows the flight path that was analysed in B and C. (B) Intensity trace showing the mean pixel value within the blue circle across each frame. The two blue insets show a single flash along with a calculated Michelson contrast. (C) Fast Fourier transform periodogram showing the relative spectral power, with the wing beat frequency labeled in red. Note: the dipterans were swarming out of reach, preventing capture and identification.

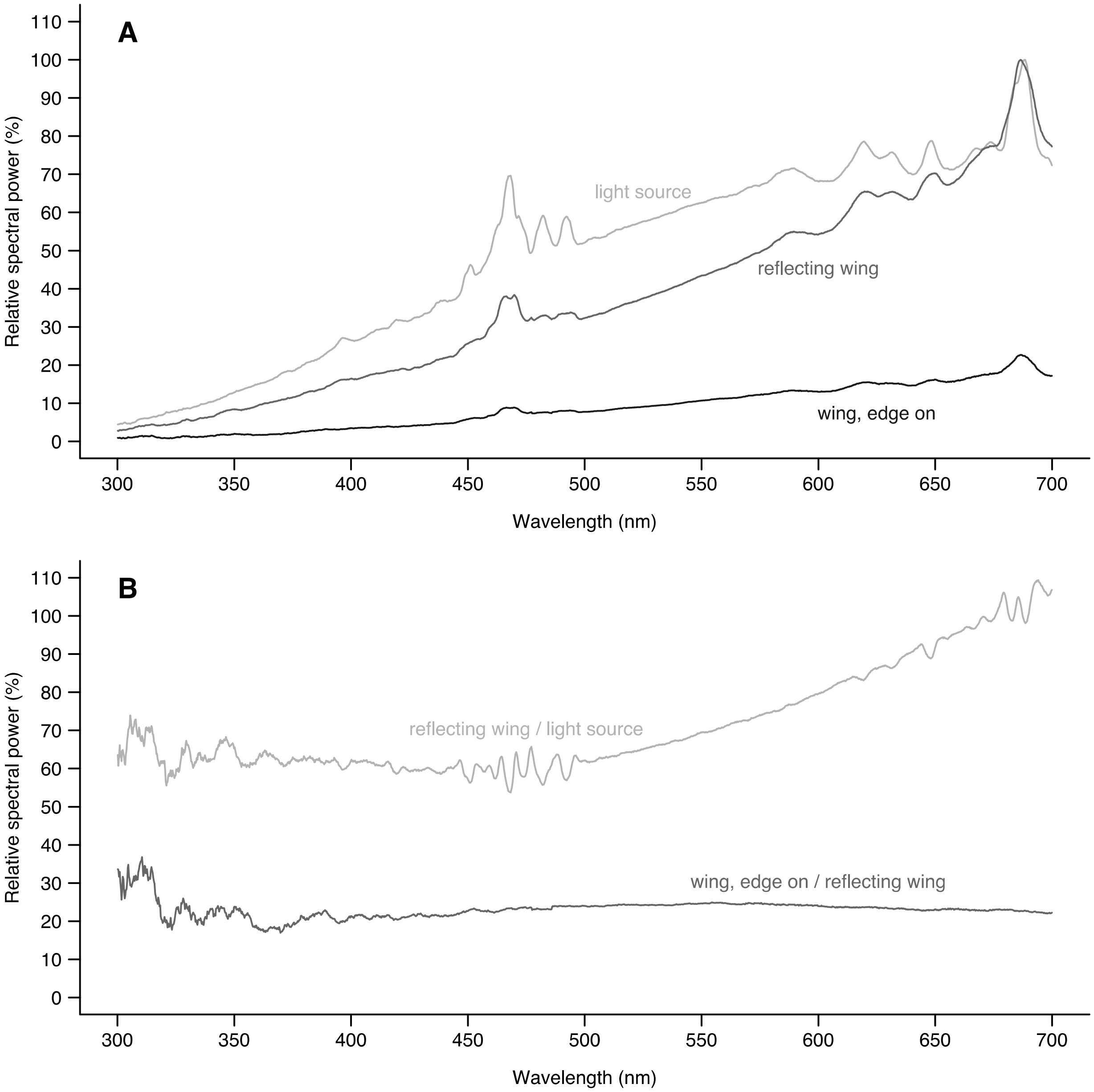


**Figure S2. Reflections from isolated single wings of *Aedes aeygpti* males**. (A) The relative spectral power of the xenon light source (as measured by the reflection of an aluminum foil square), the specularly reflecting male wing, and the same wing measured edge on. The light source and reflecting wing are normalized to their peak, whereas the edge on wing is normalized to the peak of the reflecting wing. (B) The spectral power of the reflecting wing relative to the light source, and the spectral power of the edge on wing relative to the reflecting wing.

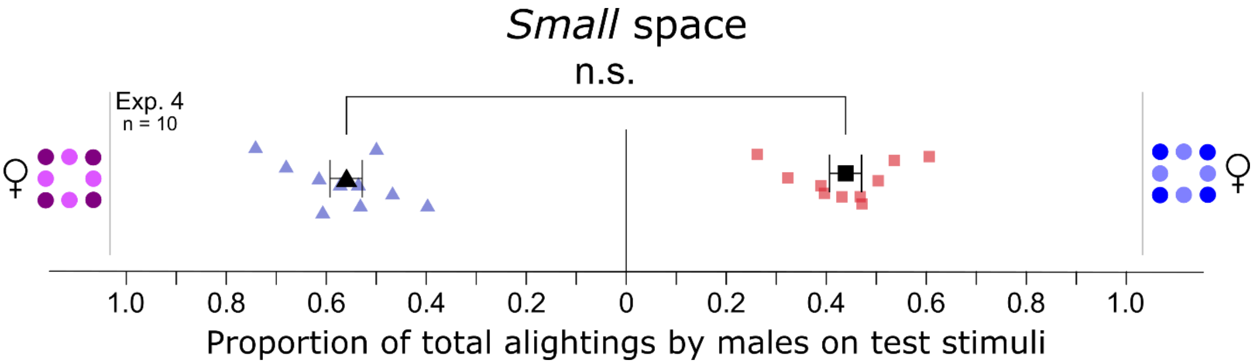


**Figure S3. Effect of wavelength on alighting responses of 2- to 7-day old male *Aedes aegypti*.** The eight purple and eight blue dots represent the number of LEDs contained within each of two LED arrays (Fig. 2B), one of which was flashing UV light and the other blue light at the 665 Hz wingbeat frequency of female *Ae. aegypti*. Each replicate was run with 50 males. Light blue triangles and light red squares show the data of individual replicates and black symbols the mean (± SE). There was no preference for either set of test stimuli (binary logistic regression model; p > 0.05; n. s. = not significant).

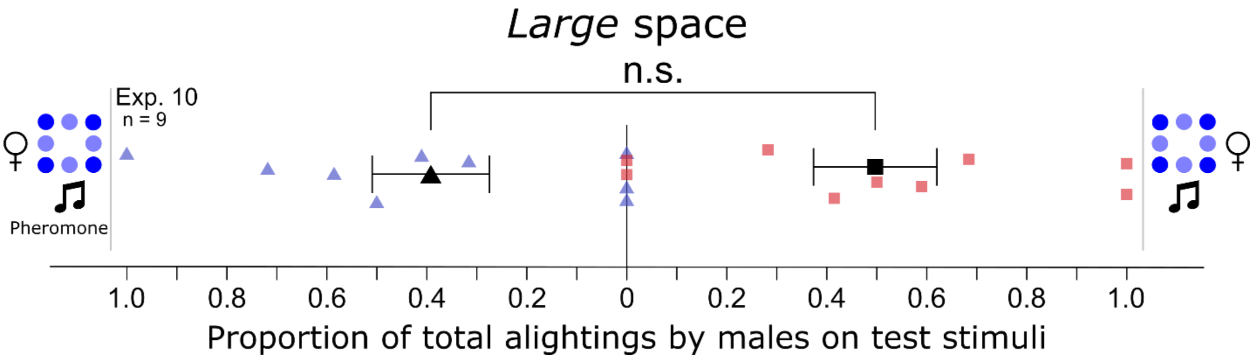


**Figure S4. Effect of ketoisophorone on the alighting responses of 2- to 7- day old male *Aedes aegypti*.** The number of blue dots represents the number of blue LEDs contained within each of two LED arrays (Fig. 2B), with LEDs flashing light at the 665 Hz wingbeat frequency of female *Ae. aegypti*. Musical notes indicate broadcast of female wingbeat sound (665 Hz) and 'Pheromone' indicates the presence of synthetic ketoisophorone (Fig. 2I), a female produced pheromone component. Light blue triangles and light red squares show the data of individual replicates and black symbols the mean (± SE). There was no preference for either set of test stimuli (binary logistic regression model; $p > 0.05$; n. s. = not significant).

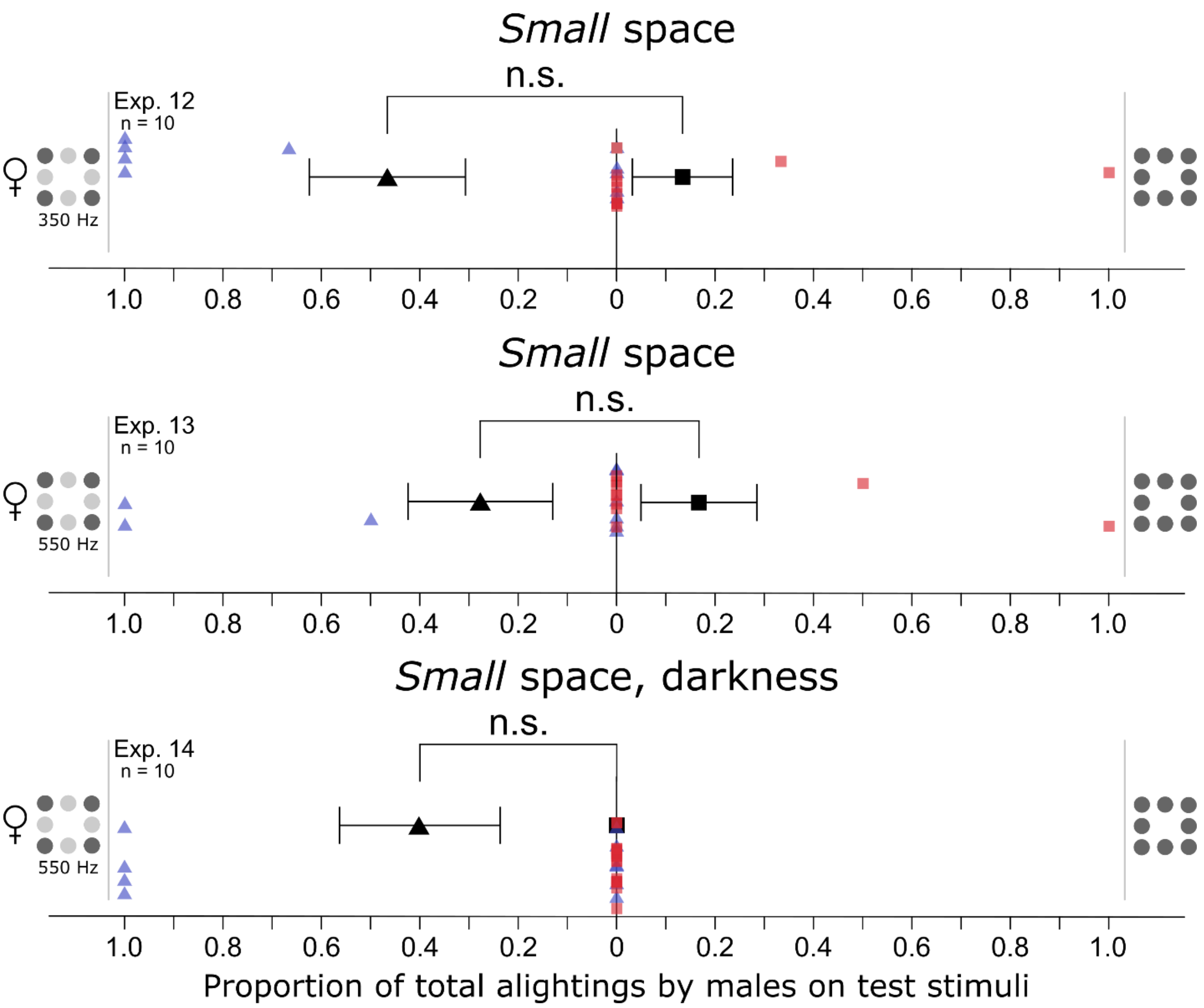


**Figure S5. Effect of visual signals on alighting responses of 2- to 7-day-old male *Culex pipiens*.** The numbers of grey dots represent the number of white LEDs contained within each of two LED arrays (Fig. 2B) that emitted either light flashes (depicted as a mixture of light- and dark-grey dots) at the reported wingbeat frequencies of *Cx. pipiens* females (350 Hz, 550 Hz) or that emitted constant light (depicted as uniformly dark-grey dots). Experiments 12 and 13 were run under standard illumination (see methods for detail), whereas experiment 14 was run at low light intensity (1 lux) with dimmed LEDs ($6.67e^{12}$ photons/$cm^2$/s). The few alighting responses (<10 total responders per replicate) recorded in these experiments did not warrant statistical analyses of data.

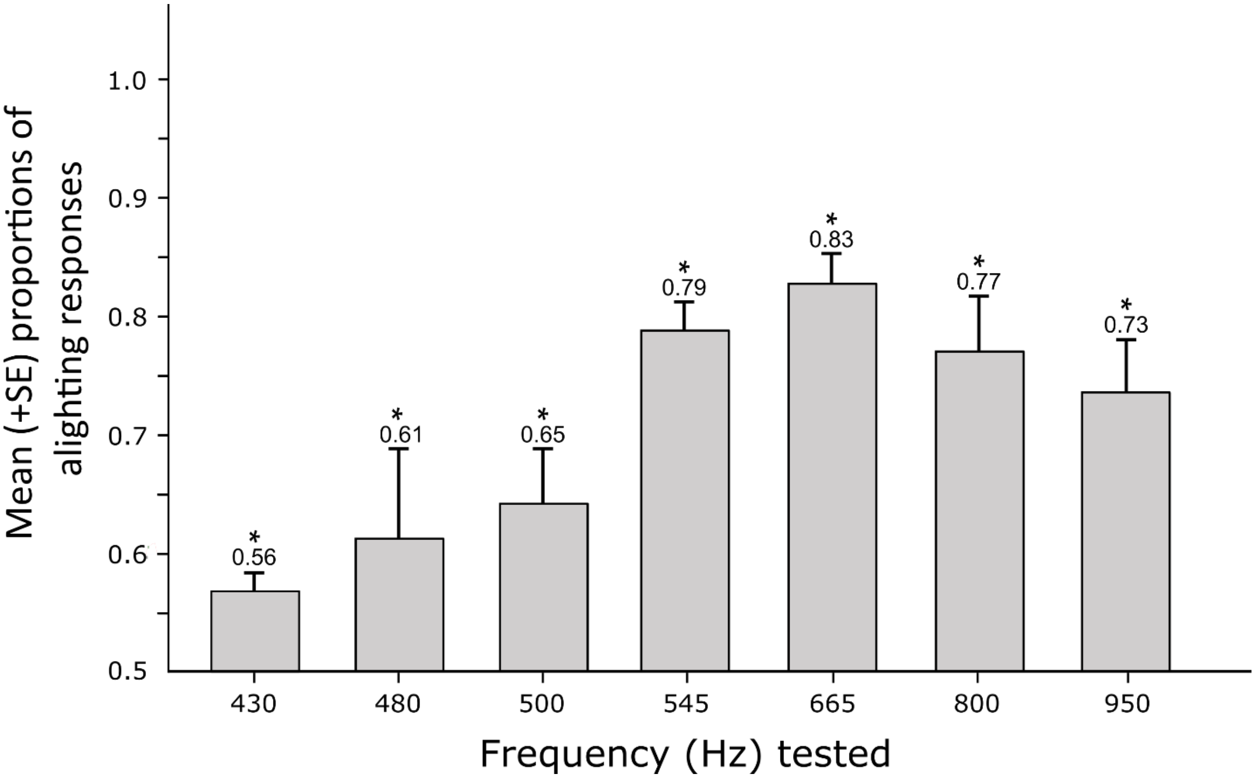


**Figure S6. Preferential alighting by *Aedes aegypti* males on LED arrays flashing light at various frequencies.** For each frequency tested, males were given a choice between two 8-LED arrays (Fig. 2B), one of which flashing white light at 430, 480, 500, 545, 665, 800 or 959 Hz, the other emitting constant white light. Proportional alighting responses are shown only for the flashing-light LED arrays. For each experiment, the asterisk indicates a significant preference for the flashing-light LED array over the constant-light LED array (*t*-test; $p < 0.05$). Figure adapted from Gries et al. (2017).

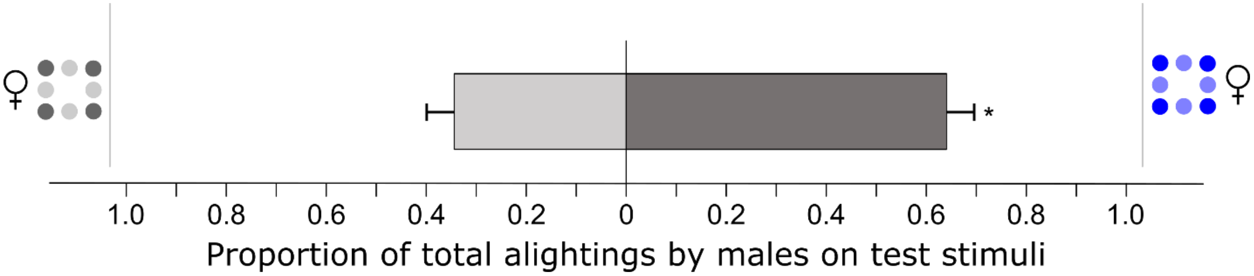


**Figure S7. Preferential alighting by *Aedes aegypti* males on LED arrays (see Figure 2B) flashing either blue or white light at 665 Hz.** The asterisk indicates a significant preference (*t*-test; $p < 0.05$). Figure adapted from Gries et al. (2017).

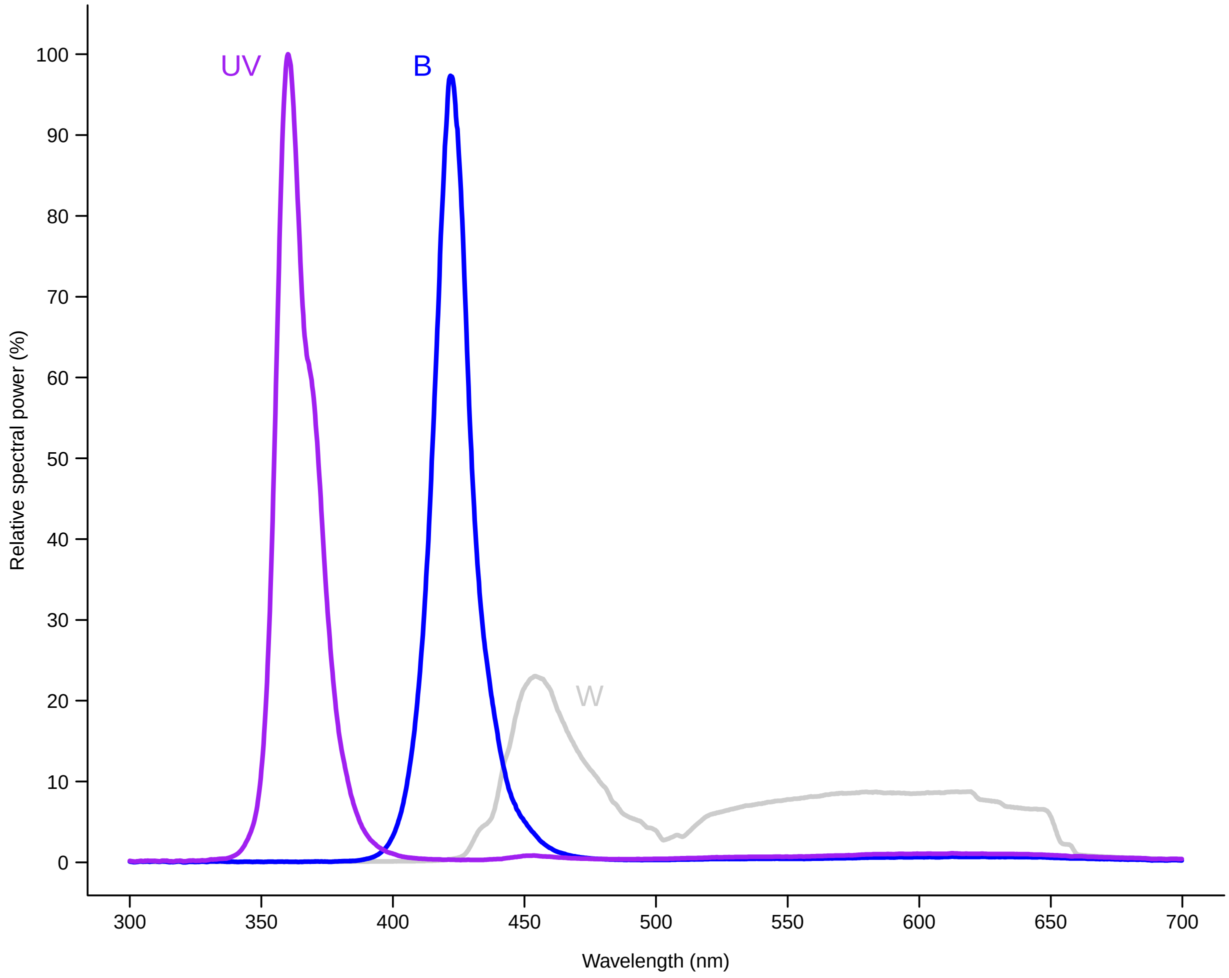


**Figure S8. Spectra of LEDs tested in behavioural bioassays.** Ultraviolet (UV), blue (B) and white (W) LEDs had intensity peaks at 360 nm, 422 nm, and at 455 nm and 620 nm, respectively. Each LED was standardized to a relative intensity of $2e^{15}$ photons/$cm^2$/s/nm.

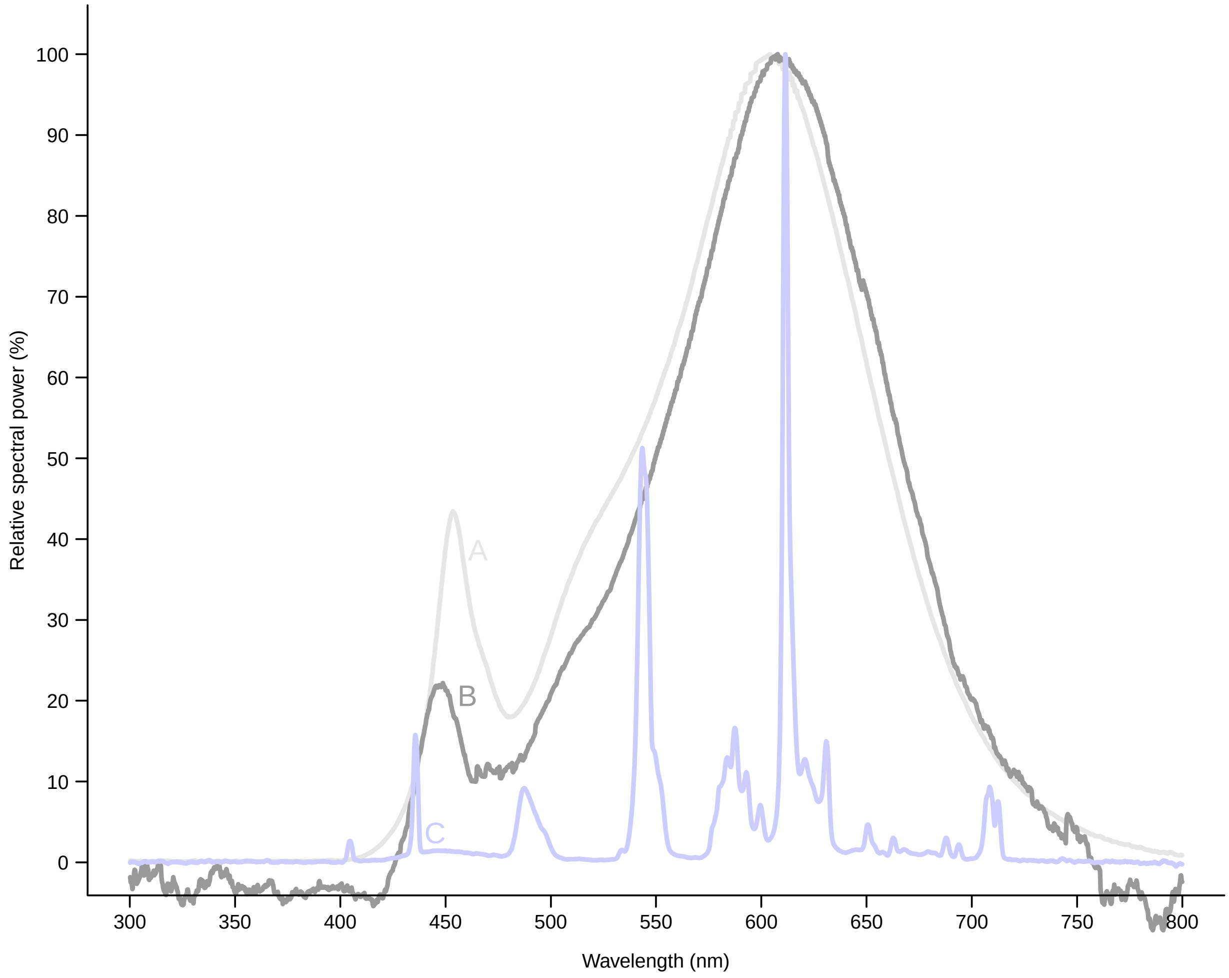


**Figure S9. Spectra of illumination used during bioassays.** Cage bioassays were illuminated with an LED bulb (Feit Electric) at full power (A) or dimmed (B). Light fixtures in the room contained two fluorescent tubes (C; General Electric). Each spectrum was normalized to it's peak.

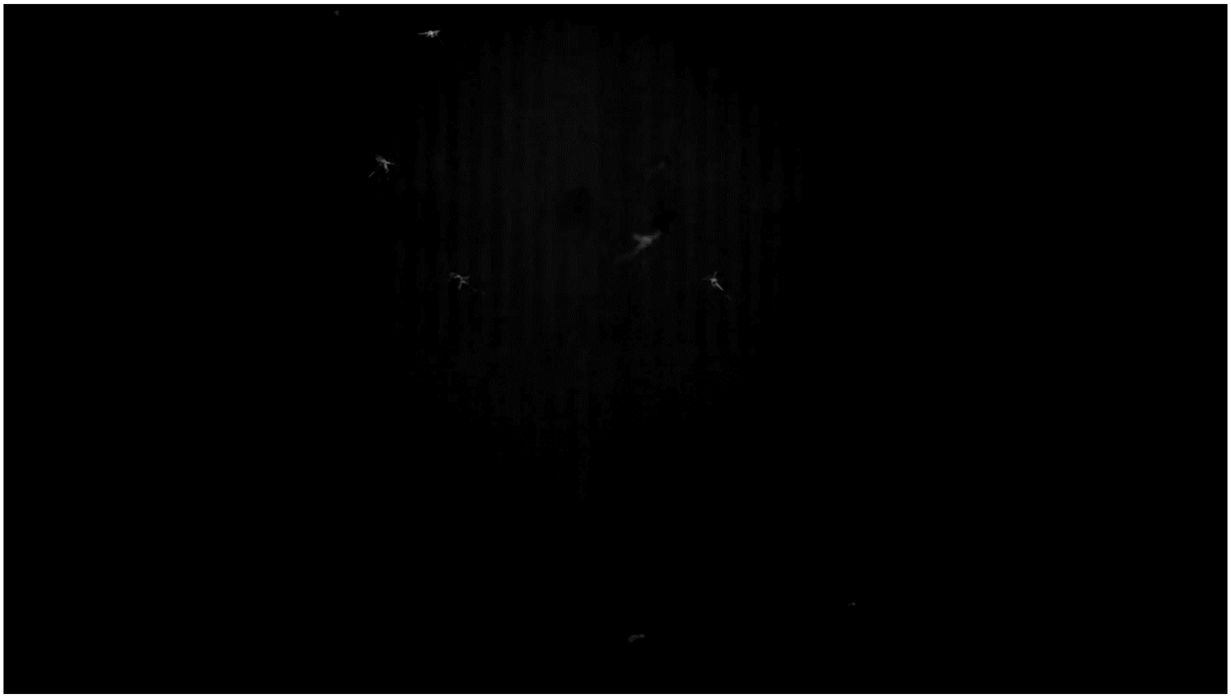

**Video S1**.[66] **Males of *Aedes aegypti* swarming in a cage in a laboratory setting.** Light flashes reflected from the wings of in-flight males are clearly visible. The video was recorded with a Photron FASTCAM NOVA S16 high-speed camera fitted with a Nikon NIKKOR Telephoto Zoom lens (AF 35-80 mm, f/4-5.6D) at a frame rate of 10000 fps and a shutter speed of 1/20000 s. Contrast and frequency analyses of the wing flash series produced by a single male from this video are shown in Fig. 1.

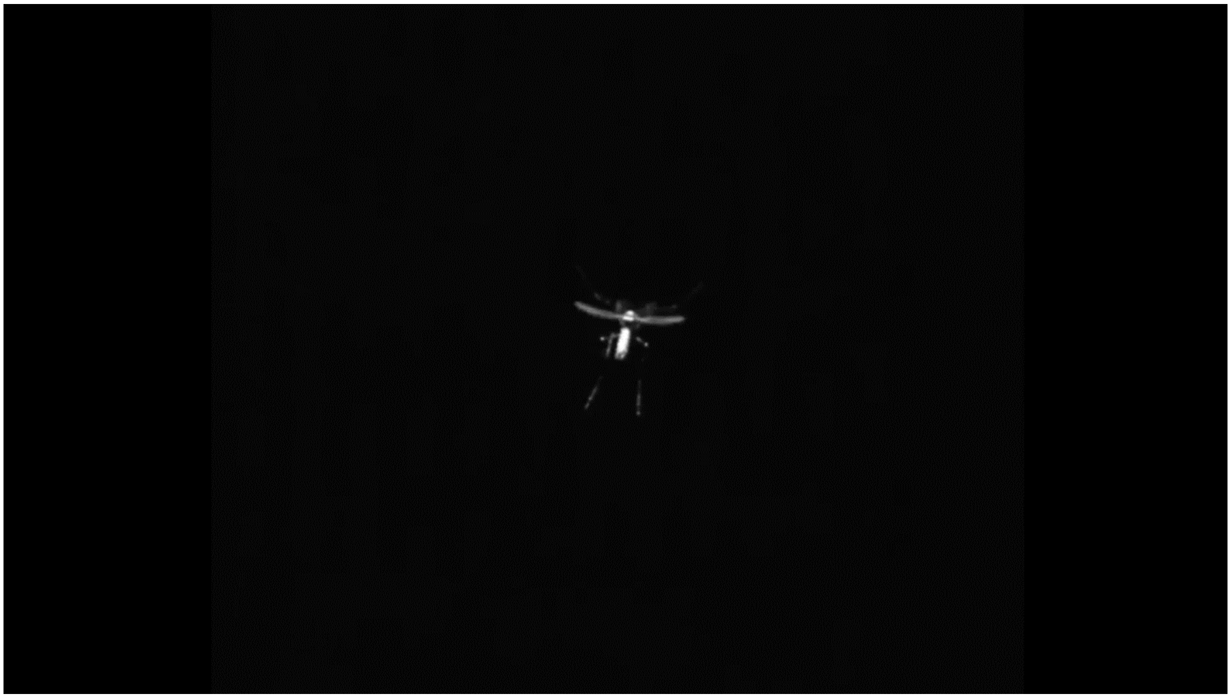

**Video S2**.[66] **Close-up of a swarming *Aedes aegypti* male.** Wing flashes appear with each flap of the male's wings. The video was recorded with a Photron FASTCAM NOVA S16 high-speed camera fitted with a Nikon NIKKOR AF-S Micro lens (105 mm, f/2.8 AF) at a frame rate of 3000 fps and a shutter speed of 1/6000 s.

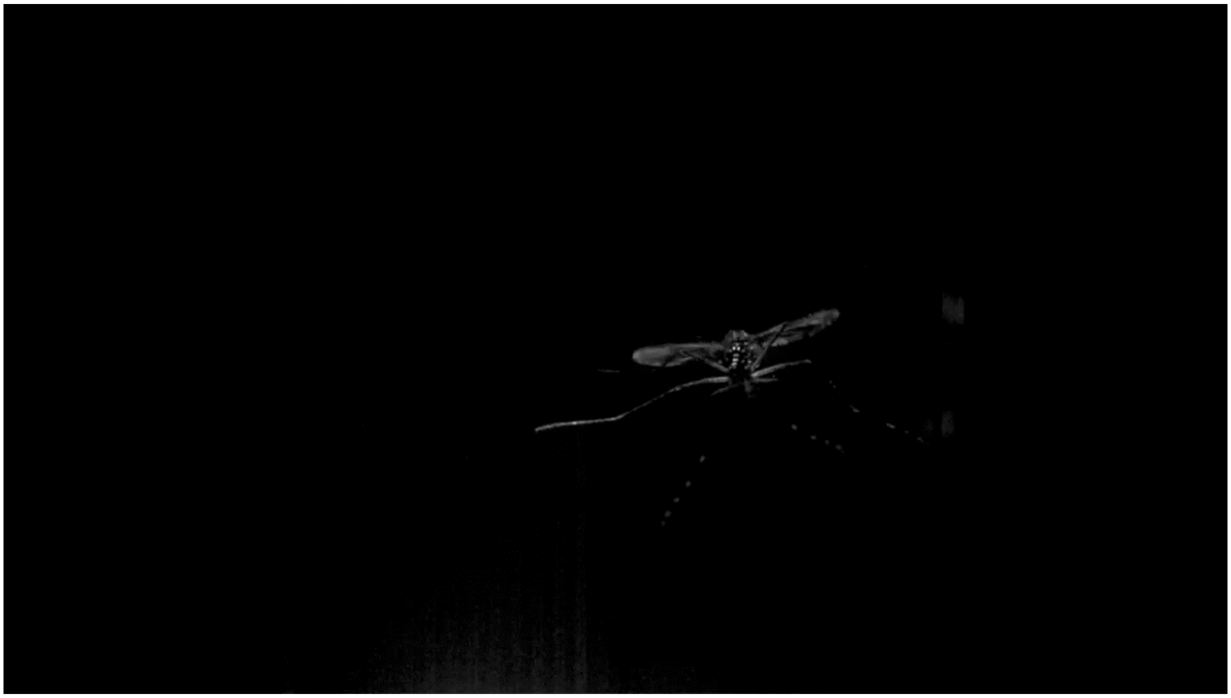

**Video S3**.[66] **Close-up of swarming *Aedes aegypti* males.** The male flying towards the camera exhibits the 'seizing and clasping' response typical of a male that approaches a female in response to her wingbeat sound.  The video was recorded with a Photron FASTCAM NOVA S16 high-speed camera fitted with a Nikon NIKKOR AF-S Micro lens (105 mm, f/2.8 AF) and a clip-on macro filter at a frame rate of 5000 fps and a shutter speed of 1/10000 s.

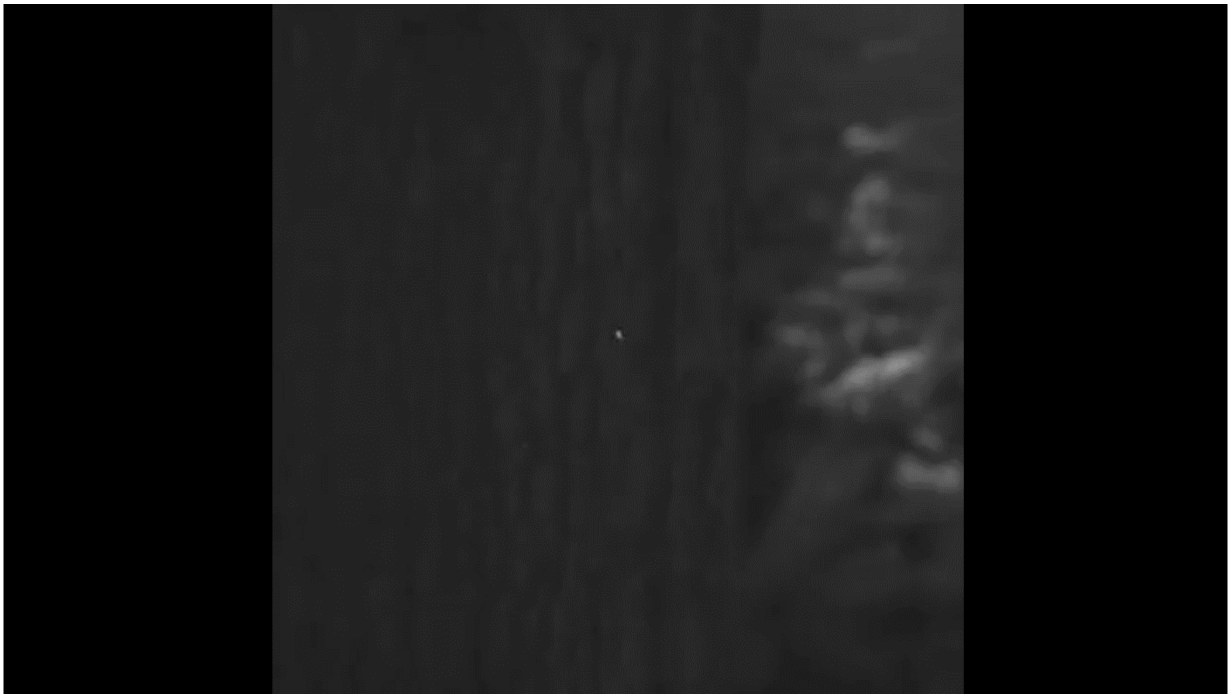

**Video S4**.[66] **Wide view of swarming dipterans in an outdoor (courtyard) setting.** These dipterans exhibited swarming behaviour with visible wing flashes under sunlight. Contrast and frequency analyses of the wing flash series produced by a single insect from this video are shown in Fig. S1. The video was recorded with a Photron FASTCAM NOVA S16 high-speed camera fitted with a Nikon NIKKOR Telephoto Zoom lens (AF 35-80 mm, f/4-5.6D) at a frame rate of 10000 fps and a shutter speed of 1/30000 s.

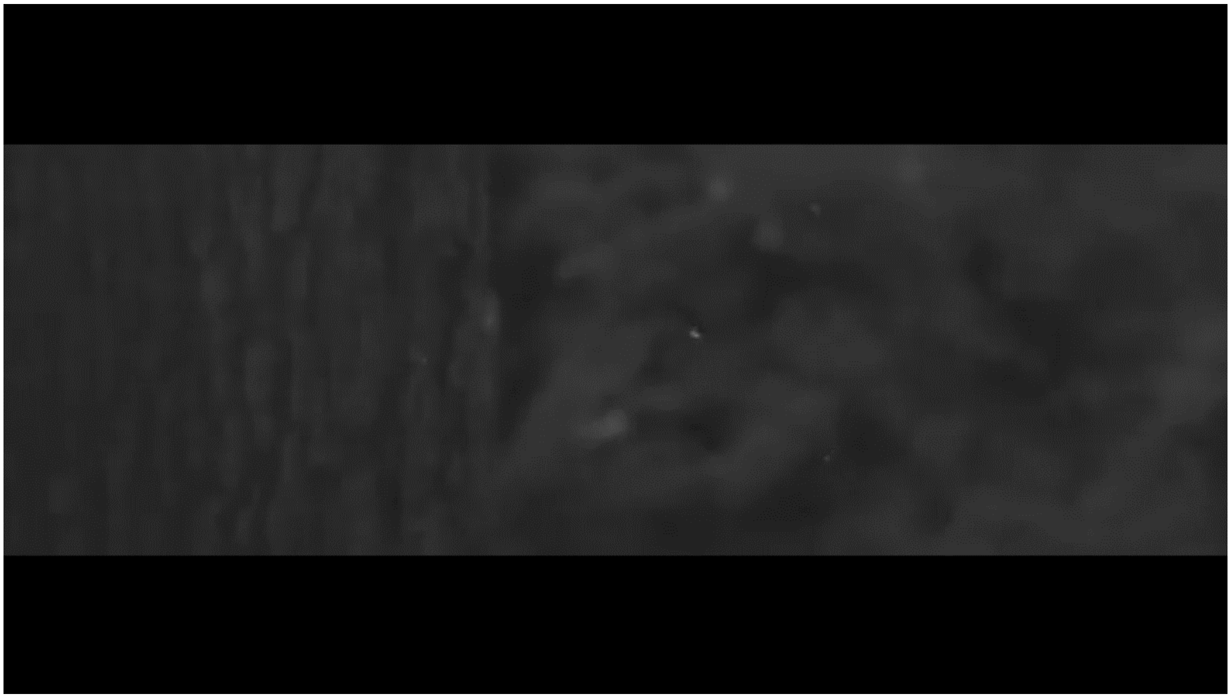

**Video S5.**[66] **Wide view of swarming dipterans in an outdoor (courtyard) setting.** These dipterans exhibited swarming behaviour with visible wing flashes under sunlight. The video was recorded with a Photron FASTCAM NOVA S16 high-speed camera fitted with a Nikon NIKKOR Telephoto Zoom lens (AF 35-80 mm, f/4-5.6D) at a frame rate of 10000 fps and a shutter speed of 1/30000 s.

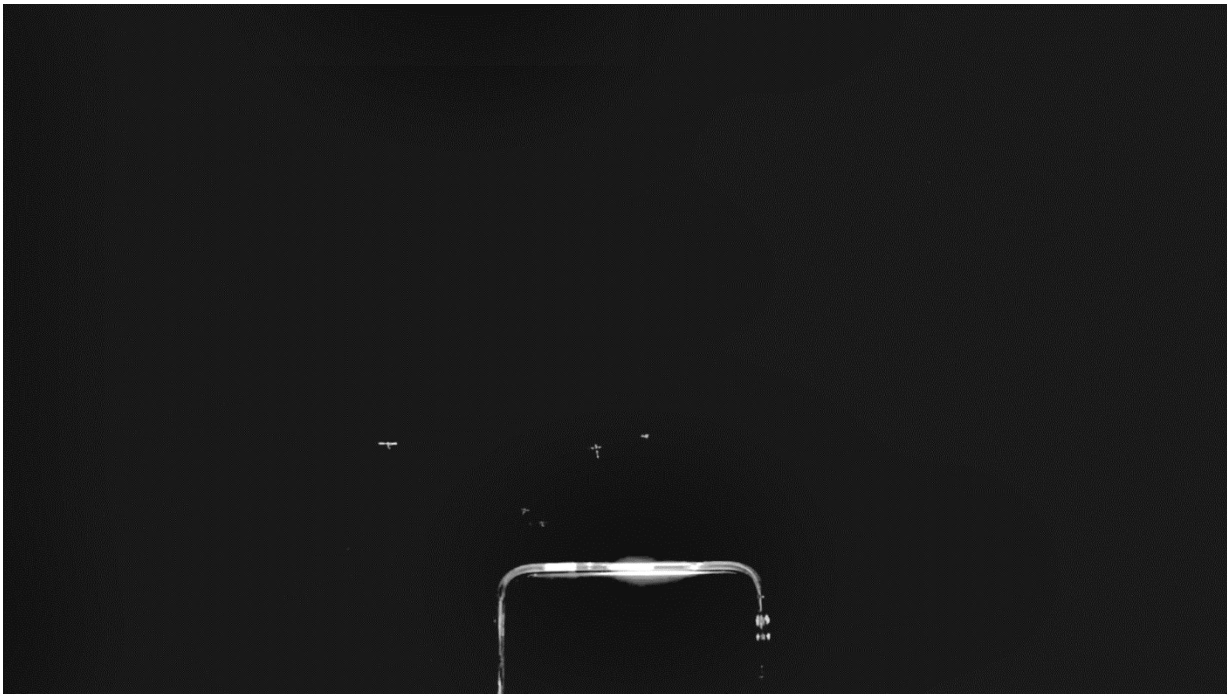

**Video S6.**[66] **Wide view of *Aedes aegypti* males swarming in a cage in a laboratory setting.** The males are responding to a 665 Hz tone played back from a smartphone in the lower portion of the frame. The males dart towards the phone’s speaker, and exhibit wing flashes under artificial light. The video was recorded with a Photron FASTCAM NOVA S16 high-speed camera fitted with a Nikon NIKKOR Telephoto Zoom lens (AF 35-80 mm, f/4-5.6D) at a frame rate of 5000 fps and a shutter speed of 1/10000 s.
